# Transcriptional Regulation of Neuropeptide Receptors Decodes Complexity of Peptidergic Modulation of Behavior and Physiology

**DOI:** 10.1101/2024.11.23.624967

**Authors:** SeungHeui Ryu, Yanan Wei, Zekun Wu, DoHoon Lee, Woo Jae Kim

## Abstract

The modulation of complex behaviors in response to environmental and physiological contexts is a fundamental aspect of animal biology, with neuropeptides (NPs) playing a crucial role in this process. This study investigates the transcriptional regulation of neuropeptide receptors (NPRs) as a mechanism for context-dependent neuropeptidergic modulation of physiology and behavior. We hypothesize that the transcriptional control of NPR genes, rather than the NPs themselves, is a critical determinant in the context-dependent modulation of behavior and physiology. Using a multi-faceted approach, including comparative genomics, transcription factor network analysis, and empirical validation in model organisms such as *Drosophila melanogaster*, we reveal a complex regulatory landscape where NPR expression is tightly controlled. Our findings demonstrate that NPR genes exhibit a higher number of enhancers, CTCF-binding sites, and open chromatin regions compared to NP genes, suggesting a greater susceptibility to transcriptional modulation. This regulatory architecture allows for precise control over neuropeptidergic signaling, enabling dynamic and context-specific behavioral and physiological responses. Our results highlight the importance of NPR-expressing cells by transcriptional regulation in mediating the effects of NPs on behavior and physiology. We show that this regulation is conserved across species, indicating an evolutionarily significant mechanism for fine-tuning neuropeptidergic signaling. Furthermore, our study provides insights into the distinct regulatory mechanisms underlying the multifunctionality of NPs and their receptors, offering a novel perspective on the transcriptional control of complex behaviors. In conclusion, this study advances our understanding of neuropeptidergic signaling by focusing on the transcriptional regulation of NPRs. Our findings have broad implications for the development of therapeutic strategies targeting neuropeptidergic systems in various neurological and behavioral disorders.

## INTRODUCTION

### Deciphering the Regulatory Mechanisms of Neuropeptide Multifunctionality

The plasticity of animal behavior is contingent upon the organism’s capacity to assimilate extrinsic and intrinsic signals from a dynamic environment, thereby regulating neuronal activity within synaptic networks of the brain. The context-specific modulation of neural circuits is predominantly mediated by non-synaptic communication involving neuropeptides (Nässel and Zandawala 2022). Neuropeptides, a diverse class of signaling molecules, play pivotal roles in modulating complex physiological processes and behaviors across various organisms, including humans and vertebrates (Hökfelt et al. 2000). These molecules are involved in a plethora of functions, such as feeding, reproduction, metabolism, and pain perception. The physiological effects of neuropeptides are primarily mediated through their interaction with specific NPRs, which are often members of the G-protein coupled receptor (GPCR) superfamily (Hoyle 1999). The intricate interplay between neuropeptides and their receptors constitutes a sophisticated signaling network that is crucial for maintaining homeostasis and adapting to environmental changes (Nässel and Winther 2010; Caers et al. 2012a; Grimmelikhuijzen and Hauser 2012; Jékely 2013; Elphick et al. 2018).

A central question in the field of neuropeptidergic signaling pertains to the mechanisms by which a single NP can induce a spectrum of behavioral and physiological outcomes, and the underlying principles that regulate its multifaceted functionalities. The multifunctional nature of neuropeptides in eliciting different behaviors and physiological responses can be attributed to a combination of factors (Dawson 1999). First, ‘receptor specificity and heterogeneity’, which describes the same neuropeptide may interact with different receptor subtypes or receptor complexes on various target cells, leading to distinct cellular responses. The diversity of NPRs and their expression patterns across different neural circuits and tissues underlies the multifunctionality of NPs. Second, ‘plurichemical transmission’, which describes that the co-release of multiple chemical messengers, including neuropeptides, with different temporal dynamics and affinities for their receptors, contributes to the varied outcomes. The interaction between NPs and other neurotransmitters (NTs) or hormones in the synaptic cleft can differentially modulate postsynaptic responses (Furness et al. 1989; Furness et al. 1995). Third, ‘cellular context and local environment’, which describes that the local environment, including the presence of enzymes, transporters, and other signaling molecules, may differentially modulate the effects of a NP. The cellular context in which a neuropeptide is released can alter its bioavailability, degradation rate, and interaction with its receptors, thus affecting its functional outcome (Nusbaum and Blitz 2012; Zhang et al. 2021). Fourth, ‘developmental and experience-dependent plasticity’, which describes that the ontogenetic timing and experience-dependent changes in the sensitivity and expression of NPs and their NPRs can lead to different behavioral and physiological responses to the different environmental context.

Within the spectrum of mechanisms proposed to elucidate the multifunctional roles of NPs, our investigation centers on the experience-dependent alterations in NPs and their corresponding NPRs, which are poised to play a pivotal role in the fine regulation of behavioral outcomes mediated by neuropeptidergic signaling (Nusbaum and Blitz 2012). The intricate web of neuropeptidergic signaling is pivotal in the orchestration of a myriad of physiological and behavioral processes across vertebrates and invertebrates (Morley 1987; van den Pol 2012; Jékely et al. 2018; Casello et al. 2022; Nässel and Zandawala 2022). NPs, as key mediators of this signaling, have been the subject of extensive research, particularly in the context of their gene expression and synthesis (Goodman 1990; Schneider et al. 1993; Minth-Worby 1994; Quinn et al. 1995; Gauthier and Hewes 2006; Romanos et al. 2017). Traditionally, studies have focused on the transcriptional regulation of NP genes as a means to understand their multifunctional roles in the nervous system. While significant insights have been gained into the mechanisms governing NP gene expression, this approach has limitations in fully elucidating the diverse and context-dependent functions of NPs.

The multifunctional nature of NPs, which enables them to modulate a wide range of physiological and behavioral responses, remains somewhat enigmatic when considered solely through the lens of peptide expression. This complexity is underscored by the fact that a single NP can elicit distinct effects in different neural circuits and under varying environmental conditions. To address this gap in knowledge, we propose a novel hypothesis that shifts the focus from the NPs themselves to the transcriptional regulation of their NPRs (Hewes and Taghert 2001; Caers et al. 2012a; Frooninckx et al. 2012; Audsley and Down 2015; Sahbaz and Iyison 2018; Beets et al. 2023; Vaudry et al. 2023). Our hypothesis posits that the transcriptional control of NPR genes is a critical determinant in the context-dependent neuropeptidergic modulation of physiology and behavior.

### Exploring the Control Mechanisms of NPs in Specific Behaviors and Physiology

The regulation of receptors, rather than ligands, in fine-tuning behavioral and physiological outputs can be more efficient for several reasons. First, receptors are highly specific to certain ligands. By regulating the receptor, one can control the response to a specific signaling molecule without affecting others. This specificity allows for more targeted adjustments in cellular responses. Second, once a ligand binds to a receptor, it can trigger a cascade of intracellular events that amplify the signal. By controlling receptor density or activity, cells can fine-tune the amplification of the signal, making the regulatory process more efficient. Third, receptors can be regulated at multiple levels (expression, trafficking, phosphorylation, etc.), providing a wide dynamic range for signal modulation. This allows for both subtle and dramatic changes in response to a constant ligand concentration. Fourth, cells can quickly adapt to changing environments by regulating receptor sensitivity or availability. For example, receptor desensitization can prevent overstimulation, while upregulation can enhance sensitivity when needed. Fifth, many ligands are essential for multiple physiological processes and are often not exclusive to a single signaling pathway. By regulating receptors, the body can conserve the ligands for use in different contexts without needing to adjust their production, which could be metabolically costly. Sixth, receptor regulation can be very rapid, allowing for quick adjustments to stimuli. This temporal control is crucial for processes that require immediate responses. Seventh, receptors can be selectively expressed or internalized in specific tissues or cell types, allowing for spatial regulation of signaling.

The dynamic regulation of neurotransmitter receptors (NTRs) is essential for maintaining the intricate balance of neuronal signaling within the central nervous system. Transcription factors (TFs) and signaling pathways activated by NT agonists play a critical role in modulating the expression levels of these receptors, thereby influencing synaptic strength and plasticity (Hadcock and Maibon 1991; Ginty et al. 1992; Lauder 1993; Lee and Masson 1993; Martin and Magistretti 1998; Desvergne et al. 2006; Albert 2012; Martinez-Lozada et al. 2016). The insulin receptor (IR) plays a central role in the regulation of glucose homeostasis and metabolism. Its proper function is crucial for maintaining systemic health, as it mediates the effects of insulin, a hormone pivotal to cellular glucose uptake and energy storage (Youngren 2007). The expression of IRs is tightly regulated at the transcriptional level, ensuring that receptor levels are adjusted in response to the body’s metabolic needs. Transcriptional regulation of the IR genes involves a complex interplay of TFs, enhancers, and epigenetic modifications that dictate the spatial and temporal expression patterns of the IRs. These regulatory elements are responsive to a variety of signals, including nutritional status, hormonal cues, and stress responses, which in turn influence the transcription of the IR genes (Cameron et al. 1992; McKeon 1994; Puig and Tjian 2005).

While the regulatory influence of receptors within the NT and hormone domains have been extensively studied, the transcriptional control exerted by neuropeptides on gene expression remains less explored. Among the neuropeptides, mammalian neuropeptide Y (NPY) stands out as a well-investigated example, particularly in terms of its interaction with its cognate receptors (Loh et al. 2015). NPY, a 36-amino acid peptide, functions as both a neuropeptide and neuromodulator across the central and peripheral nervous systems. Upon binding to its GPCRs (Y1, Y2, Y4, Y5), NPY triggers a series of intracellular signaling cascades that are pivotal for modulating gene expression. The transcriptional effects of NPY are well-documented, with evidence indicating its capacity to regulate the expression levels of its own receptors, as well as genes integral to neuronal function and survival (MINTH and DIXON 1990; Minth-Worby 1994; Magni 2003; Eva et al. 2006). For example, the engagement of NPY receptors can activate downstream signaling molecules, such as extracellular signal-regulated kinases (ERK) and protein kinase A (PKA). These molecules, in turn, are capable of phosphorylating and thereby modulating the activity of key transcription factors, including the cAMP response element-binding protein (CREB). This intricate regulatory network underscores the significance of NPY in the transcriptional landscape of neuronal cells (Sabol and Higuchi 1990; Minth-Worby 1994; Lerchen et al. 1995; Herzog et al. 1997; Sheriff et al. 1998; Thorsell et al. 1999; Higuchi et al. 2005; Eva et al. 2006; He et al. 2024).

### Advantages of Invertebrate Models in Unraveling NPRs’ Transcriptional Regulation

The exploration of NPs began with significant discoveries in mammals, notably the hypothalamic neuropeptides vasopressin and oxytocin (Donaldson and Young 2008). These peptides have a dual role, acting as hormones to regulate essential functions like diuresis and lactation, and influencing social behaviors within the brain. This foundational work laid the groundwork for understanding NPs, but it was the study of insects that truly expanded our knowledge. Insects, with their relatively simple nervous systems, have proven to be excellent models for investigating the intricacies of NP-NPR systems (Nässel 1996; Nässel and Homberg 2006; Nässel et al. 2008; Nässel 2009; Nässel and Winther 2010; Nässel and Wegener 2011; Kapan et al. 2012; Nässel et al. 2019; Nässel and Zandawala 2019; Zandawala et al. 2021; Nässel and Zandawala 2022).

NPs and NPRs have an early evolutionary origin and are already abundant in basal animals with primitive nervous systems such as cnidarians (Hydra, jellyfishes, corals, and sea anemones) (Grimmelikhuijzen and Hauser 2012). This evolutionary antiquity underscores the fundamental role of NPs in diverse organisms. The molecular identification of NPs in invertebrates, such as the adipokinetic hormone (AKH) and proctolin in insects, and the cardioexcitatory neuropeptide FMRFamide in mollusks, has further cemented the importance of these molecules in biological processes. These findings in insects, which are among the first animals where NPs were characterized, highlight their suitability for NP research (Brown 1975; Starratt and Brown 1975; Stone et al. 1976; Price and Greenberg 1977). The study of NPs in insects not only provides insights into their basic biology but also offers a window into the evolution of these signaling molecules. As we investigate deeper into the NP landscape, insects continue to serve as valuable models, offering a clear view of the foundational mechanisms that have persisted throughout the evolution of more complex organisms (Nässel and Winther 2010; Schoofs et al. 2016; Nässel and Zandawala 2022).

*Drosophila melanogaster* offers a unique and valuable model for studying the transcriptional regulation of NPRs due to several compelling reasons. Firstly, the evolutionary conservation of neuropeptide signaling pathways across invertebrates and vertebrates allows researchers to investigate fundamental mechanisms that are likely to be preserved in more complex organisms (Hoyle 1999; Caers et al. 2012a; Grimmelikhuijzen and Hauser 2012; Jékely 2013; Elphick et al. 2018). Secondly, *D. melanogaster* has been extensively studied, and a wealth of genetic resources, including mutants, transgenic lines, and gene editing technologies, are readily available. These tools enable researchers to manipulate gene expression and observe the consequences in a controlled manner, providing a deeper understanding of NPR transcriptional regulation (Bellen et al. 2010; Venken et al. 2011; Yamamoto et al. 2014; Nagarkar-Jaiswal et al. 2015; Wangler et al. 2015). Moreover, the variety and accessibility of sequencing databases for *D. melanogaster* have expanded significantly in recent years. High-throughput sequencing technologies have generated vast amounts of transcriptomic data for numerous invertebrate species. This wealth of information allows for comparative analyses, identification of conserved regulatory elements, and the discovery of novel NPRs. The ease of obtaining such data facilitates hypothesis generation and the design of targeted experiments to explore the transcriptional regulation of NPRs (Bargmann 2012; Maisak et al. 2013; Ito et al. 2014; Takemura 2014; Ohyama et al. 2015; Bates et al. 2020; Scheffer et al. 2020; Li 2021; Li et al. 2022a; Court et al. 2023). Furthermore, the simplicity of some invertebrate nervous systems, compared to their vertebrate counterparts, can be advantageous for dissecting complex regulatory networks (Leadbeater and Chittka 2007; Zhu 2013; Douglas 2019; Nässel and Zandawala 2022).

By addressing above questions and testing the proposed hypothesis, we aim to uncover the mechanisms how the same neuropeptide can exert diverse effects on behavior and physiology, advancing our understanding of neural circuit function and plasticity.

## RESULT

### Differential Cis-Regulatory Landscapes of Human Neuropeptide and Neuropeptide Receptor Genes

In the cellular orchestration of gene expression, transcriptional regulation acts as the pivotal mechanism dictating the temporal and conditional activation or repression of genes within the cellular milieu. This process is akin to an intricate set of directives that delineates the precise moments and conditions under which a cell transcribes a gene’s DNA sequence into a functional RNA and protein (Bintu et al. 2005). TFs are pivotal regulatory proteins that govern this process, serving as molecular switches that modulate gene expression. These factors exhibit high specificity, binding to discrete DNA sequences termed transcription factor binding sites (TFBS). The interaction between a TF and its cognate TFBS is reminiscent of a key-lock mechanism, where each TF is uniquely tailored to engage with a particular DNA sequence (Todeschini et al. 2014).

Enhancers are pivotal regulatory elements that orchestrate gene expression patterns essential for a myriad of biological processes in multicellular organisms. Considering their multifaceted roles, it is expected that enhancers exhibit a broad spectrum of architectural variations, including differences in length, the quantity of TFBSs, and the binding specificity of the associated TFs. Typically, enhancers span approximately 10 to 1,000 base pairs (bps) and contain a few to several dozen TFBSs (Blackwood and Kadonaga 1998; Yáñez-Cuna et al. 2013). The TFBSs, located on the DNA sequence, are widely recognized as critical regulatory hubs that govern gene expression. The influence of TFBSs on their target genes is often cumulative, with each site contributing to the overall regulatory effect with varying intensities and degrees of redundancy (Spivakov 2014). There is a consensus that the length and number of an enhancer and the composition of its TFBSs—their number and specific identities—are indicative of the enhancer’s regulatory complexity. Enhancers tasked with more intricate regulatory functions tend to be longer and harbor a greater number of TFBSs, albeit with potentially lower binding specificities. The increased number of binding sites allows for a multitude of combinatorial arrangements, enabling enhancers to direct a diverse array of expression patterns and, consequently, cellular outcomes, utilizing a finite set of TFs (Li and Wunderlich 2017).

To elucidate the distinctions in transcriptional regulatory elements between neuropeptide (NP) and neuropeptide receptor (NPR) genes, we conducted a comparative analysis of their cis-regulatory DNA components across human (*Homo sapiens*), mice (*Mus musculus*), and fruit fly (*Drosophila melanogaster*) species (see “Materials and Methods” section). We observed a significant disparity in the number of enhancers associated with NP genes compared to those associated with NPR genes, irrespective of whether the NPs were unpaired or paired with their respective NPRs (Fig. 1A–B). Conversely, the number of TFBSs per gene was found to be similar between NP and NPR genes, under both unpaired and paired conditions (Fig. S1A–B), suggesting that the component of TFBSs not the number of TFBSs is crucial for transcriptional regulation for complexity as suggested by others (Li and Wunderlich 2017).

**Fig. 1.**
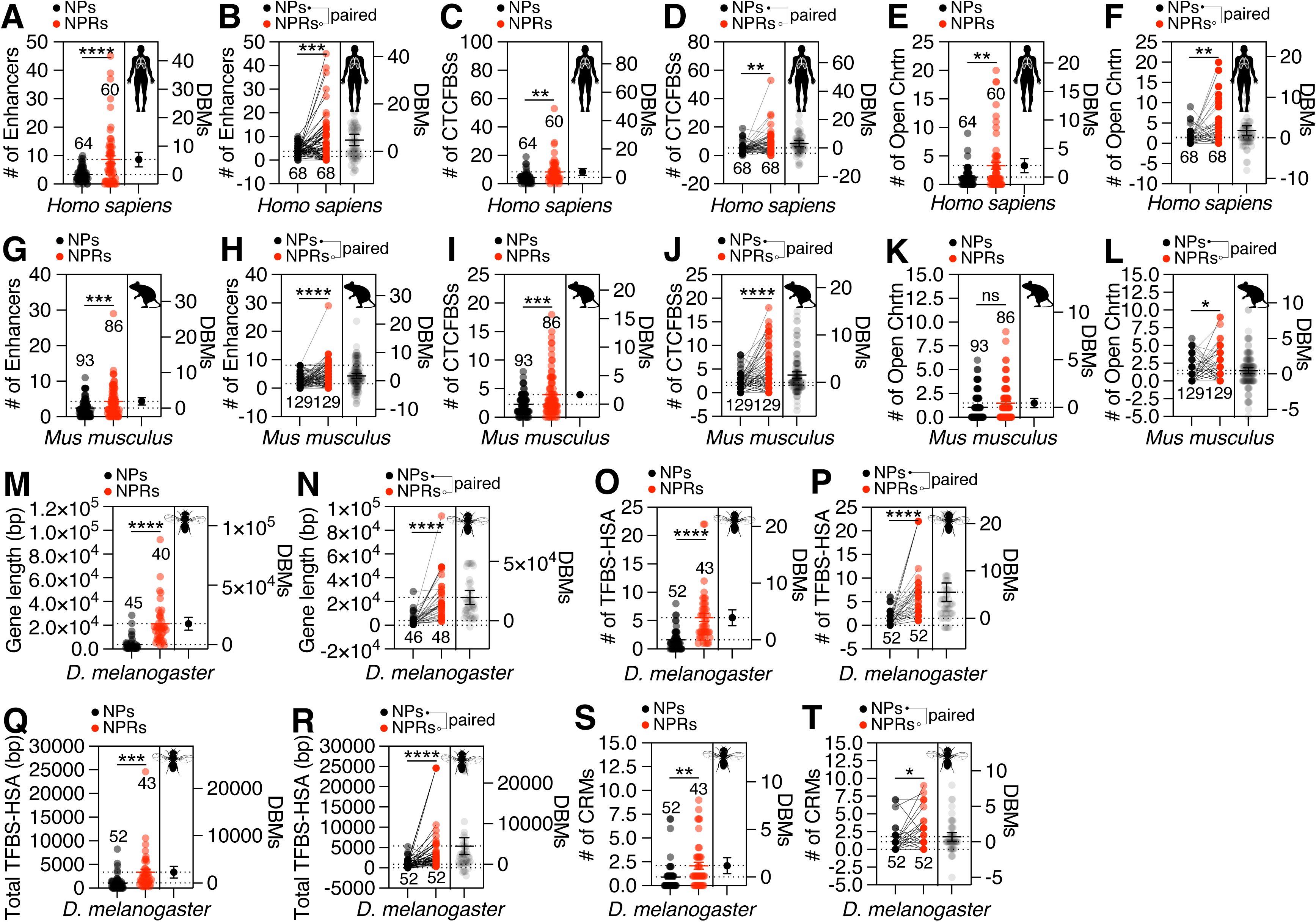
Neuropeptide receptor (NPR) Genes Exhibit a More Complex Regulatory Architecture than neuropeptide (NP) Genes Across Species. **A**–**L**. Similar Regulatory Landscapes of *Homo sapiens* and *Mus musculus*, displaying the counts of enhancers (a–b, g–h), CTCF-binding sites (CTCFBS) (c–d, i–j), and open chromatin (Chrtn) regions (e–f, k–l) associated with NP and NPR genes, along with the ‘difference between means (DBMs)’ between NP and NPR genes in *H. sapiens* (a–f) and *M. musculus* (g–l). Each dot in the plot represents the number of gene regulatory elements of each NP or NPR genes, and the solid line connects the data for the NP-NPR pairs. **M–T**. Corresponding Regulatory Landscape in *Drosophila melanogaster*, displaying the gene length (m-n), average (o-p) and total (q-r) length of transcription factor binding site hotspots (TFBS-HSA), and abundance of cis-regulatory modules (CRMs) (s-t) in NP and NPR genes, along with the DBMs between NP and NPR genes in *D. melanogaster*. The mean value and standard error are labeled within the dot plot (black lines). DBMs represent the ‘difference between means’ for the evaluation of estimation statistics (See **MATERIALS AND METHODS**). Asterisks represent significant differences, as revealed by the unpaired Student’s t test, and ns represents non-significant differences (*\*p<0.05, **p<0.01, ***p< 0.001, ****p< 0.0001*).

The CCCTC-binding factor (CTCF), an evolutionarily conserved and multifunctional transcription regulator, plays a pivotal role in mediating chromosomal interactions, ranging from interchromosomal to intrachromosomal levels. As an architectural protein, CTCF is instrumental in establishing boundaries between topologically associating domains (TADs) and in facilitating interactions between regulatory sequences within these domains. The consensus sequence for CTCF-binding sites (CTCFBSs) is highly conserved across species (Holwerda and Laat 2013; Ong and Corces 2014). Utilizing CTCFBSDB 2.0, a comprehensive database for CTCF-binding sites and genome organization (Ziebarth et al. 2012), we discerned a significant preponderance of CTCFBSs in NPR genes over NP genes (Fig. 1C–D).

Chromatin architecture is intricately linked to the regulation of gene transcription (Hübner et al. 2013). Eukaryotic genomes are organized into a dynamic hierarchy of chromatin structures, where the transition from a repressive, compact chromatin state to a more open, transcriptionally permissive state is a critical step in the transcriptional process (Li et al. 2004). Our analysis revealed a notably higher number of open chromatin regions in NPR genes compared to NP genes, in both unpaired and paired scenarios (Fig. 1E–F). This observation suggests that the genomic regions encompassing NPR genes are more prone to transcriptional regulation.

In conclusion, the collective findings from our analysis indicate that the genomic landscapes of NPR genes in human are more intricately tailored for transcriptional regulation compared to those of NP genes, highlighting the distinct regulatory mechanisms that underpin their functional roles in metazoan biology.

### Cis-Regulatory Element Disparity in Murine NP and NPR Genes

In our subsequent analysis, we examined the cis-regulatory elements of murine NPs and NPRs genes. Our findings reveal a significant disparity in the enhancer repertoire, with NPR genes harboring a greater number of enhancers compared to NP genes, a pattern congruent with the observations made in human (Fig. 1G–H). Similarly, the count of CTCFBSs was notably higher in NPR genes (Fig. 1I–J). Interestingly, an increased number of open chromatin regions was observed exclusively when NP and NPR genes were considered in paired contexts (Fig. 1K–L). Collectively, these results indicate a conserved cis-regulatory landscape between human and mouse, underscoring the importance of these regulatory elements in the transcriptional control of NP and NPR genes.

Variation in human gene lengths is predominantly due to differences in intron sizes, with longer genes containing more intronic sequences. Introns are more extensive in humans and in tissue-specific genes compared to housekeeping genes (Eisenberg and Levanon 2003). Beyond their roles in genome organization and RNA processing, intron length may modulate gene expression dynamics, particularly for genes activated by stress or cellular signals. The shorter average length of housekeeping genes compared to other genes suggests that an increase in non-housekeeping gene size has been evolutionarily advantageous (Swinburne et al. 2008). This could allow longer genes to exhibit greater expression plasticity, enabling nuanced responses to various developmental, environmental, and physiological stimuli (Kirkconnell et al. 2017). Thus, gene length, influenced by intronic content, may have functional implications for gene regulation and cellular adaptation. Strikingly, our analysis indicates that the genomic length of NPR genes is markedly greater than that of NP genes (Fig. S1C–D). This distinction in gene length implies a higher susceptibility of NPR genes to transcriptional modulation in response to temporal stimuli, relative to NP genes. This suggests that the architectural features of NPR genes may confer a greater capacity for dynamic gene expression, enabling a more responsive transcriptional regulation under varying conditions.

GO annotation, which stands for Gene Ontology annotation, is a systematic approach to functionally describe genes and their products, such as proteins, in a structured, controlled vocabulary that cover three main domains, molecular function, biological process, and cellular component (Plessis et al. 2011; Ashburner and Lewis 2016; Consortium 2019). The comparative analysis of GO annotations reveals a similar quantity of annotations for murine NP and NPR genes (Fig. S2A-D), indicating that both NP and NPR genes may be involved in a comparable level of functional complexity. It’s important to note that while the number of GO annotations might be similar, the specific annotations (i.e., the actual GO terms) might differ significantly between NP and NPR genes due to their distinct biological roles. Therefore, a detailed analysis of the specific GO terms associated with each gene type would provide a more nuanced understanding of their functional similarities or differences.

### Comparative Cis-Regulatory Landscape of NP and NPR Genes in *Drosophila melanogaster*: Implications for Transcriptional Dynamics

*Drosophila melanogaster*, commonly known as the fruit fly, has been a pioneering model in the field of transcriptional regulation, contributing seminal insights into the molecular mechanisms governing gene expression in animals. Key concepts—such as the action of cis-regulatory elements over extended genomic distances, the orchestration of developmental processes by hierarchical networks of transcription factors, the equalization of gene dosage between sexes (dosage compensation), and the variegation in gene expression due to chromosomal position effects—have been elucidated through research on this organism (Biggin and Tjian 2001; Bier 2005). The study of *Drosophila*’s transcriptional regulation has been pivotal for deepening our comprehension of the genetic and molecular underpinnings of human diseases, offering a robust framework for the development of innovative diagnostic approaches and therapeutic interventions (FitzGerald et al. 2006; Bellen et al. 2010; Spivakov et al. 2012). Mirroring the chromatin landscape of humans, active genes in *Drosophila* exhibit unique chromatin signatures that correlate with their gene length, exon-intron structure, regulatory roles, and genomic contexts, highlighting the intricate interplay between chromatin architecture and gene regulation (Kharchenko et al. 2011).

In the mouse genome, as depicted in Fig. S1C–D, NPR genes exhibit a significantly greater gene length compared to NP genes, which is also observed in *Drosophila*, implying a higher propensity for transcriptional modulation in NPR genes (Fig. 1M–N). Furthermore, the abundance and total length of transcription factor binding sites identified through hot spot analyses (TFBS-HSA) is markedly higher within the cis-regulatory regions of NPR genes than in those of NP genes, irrespective of whether they are unpaired or paired (Fig. 1O–R). Despite discernible differences in the quantity of TFBS-HSA NP and NPR genes, the mean length of these TFBS-HSA elements was found to be similar between the two gene classes (Fig. S1E–F). This observation implies that the diversity of TFBS components, rather than their linear dimensions, may exert a pivotal influence on the modulation of gene transcription. The comparable lengths of TFBS-HSA across NP and NPR genes suggest that the complexity of transcriptional regulation is more likely dictated by the repertoire of transcription factor interactions and the combinatorial arrangements of binding sites, rather than by the mere extension of the binding site sequences themselves.

Cis-regulatory modules (CRMs), which are non-coding segments of DNA, play an indispensable role in governing gene expression (Borok et al. 2009). As a collective designation for regulatory elements situated outside the core promoter region, CRMs are instrumental in modulating gene expression with precise spatio-temporal resolution (Kazemian and Halfon 2018; Rivera et al. 2019; Keränen et al. 2022). Our analysis reveals a marked increase in both the quantity and aggregate length of CRMs within NPR genes compared to NP genes (Fig. 1S–T). However, when CRMs are examined in the context of paired genes, no significant differences in the average length of CRMs are observed between NP and NPR genes (Fig. S1I–J). These findings, akin to the observations from the TFBS-HSA, indicate that the diversity and prevalence of CRMs, rather than their individual lengths, are pivotal for the transcriptional regulation of NPR genes. This underscores the importance of the composition and complexity of the cis-regulatory landscape in shaping the transcriptional output of these genes.

In the culmination of our analysis, we observed a congruent distribution of tissue expression patterns annotated in the *Drosophila melanogaster* atlas for NP and NPR genes (Fig. S1K–L). This congruence implies that the diversity of tissues in which NPs and NPRs are expressed does not exhibit a significant disparity. The observed equivalence in tissue expression profiles posits that the transcriptional regulation landscape of NPRs may have evolved to prioritize temporal modulation over spatial specificity in gene expression regulation. This inference is drawn from the premise that similar tissue expression patterns across NP and NPR genes necessitate a regulatory strategy that can accommodate dynamic and context-dependent changes in gene expression, rather than one that is strictly confined to spatial constraints.

### Post-transcriptional Regulatory Landscape of NP and NPR-Encoding Genes

Gene expression is a multifaceted process, with post-transcriptional regulation representing a pivotal stage that fine-tunes the production of functional proteins (Halbeisen et al. 2007). This level of control is essential for several reasons. Firstly, it allows cells to rapidly respond to changing environmental conditions and cellular signals by modulating the stability and translation efficiency of specific mRNAs. Secondly, post-transcriptional mechanisms enable the generation of protein diversity through processes such as alternative splicing, which can produce multiple protein isoforms from a single gene. Thirdly, it ensures the precise spatial and temporal control of protein synthesis, which is crucial for proper cellular function and the development of complex organisms (Halbeisen et al. 2007; Franks et al. 2017; Corbett 2018). The post-transcriptional regulation landscape of genes encoding NPs and NPRs is particularly important, given the critical roles these molecules play in neuronal signaling and various physiological processes (Lerchen et al. 1995).

In the context of the post-transcriptional regulation landscape, various mRNA features play significant roles in determining the fate and function of the encoded proteins. The total length of an mRNA molecule, encompassing both the coding and non-coding regions, influences its stability, translation efficiency, and subcellular localization. Longer mRNAs might be subject to more complex regulation due to increased opportunities for secondary structure formation, regulatory element interactions, and greater susceptibility to degradation (Santiago et al. 1986; Masuyama et al. 2004; Fernandes et al. 2017; Zhang et al. 2020). In both unpaired and paired gene contexts, the mRNA molecules of NPR genes exhibit a significantly greater length compared to those of NP genes in both *Murine* and *Drosophila* species (Fig. 2A–D).

**Figure 2.**
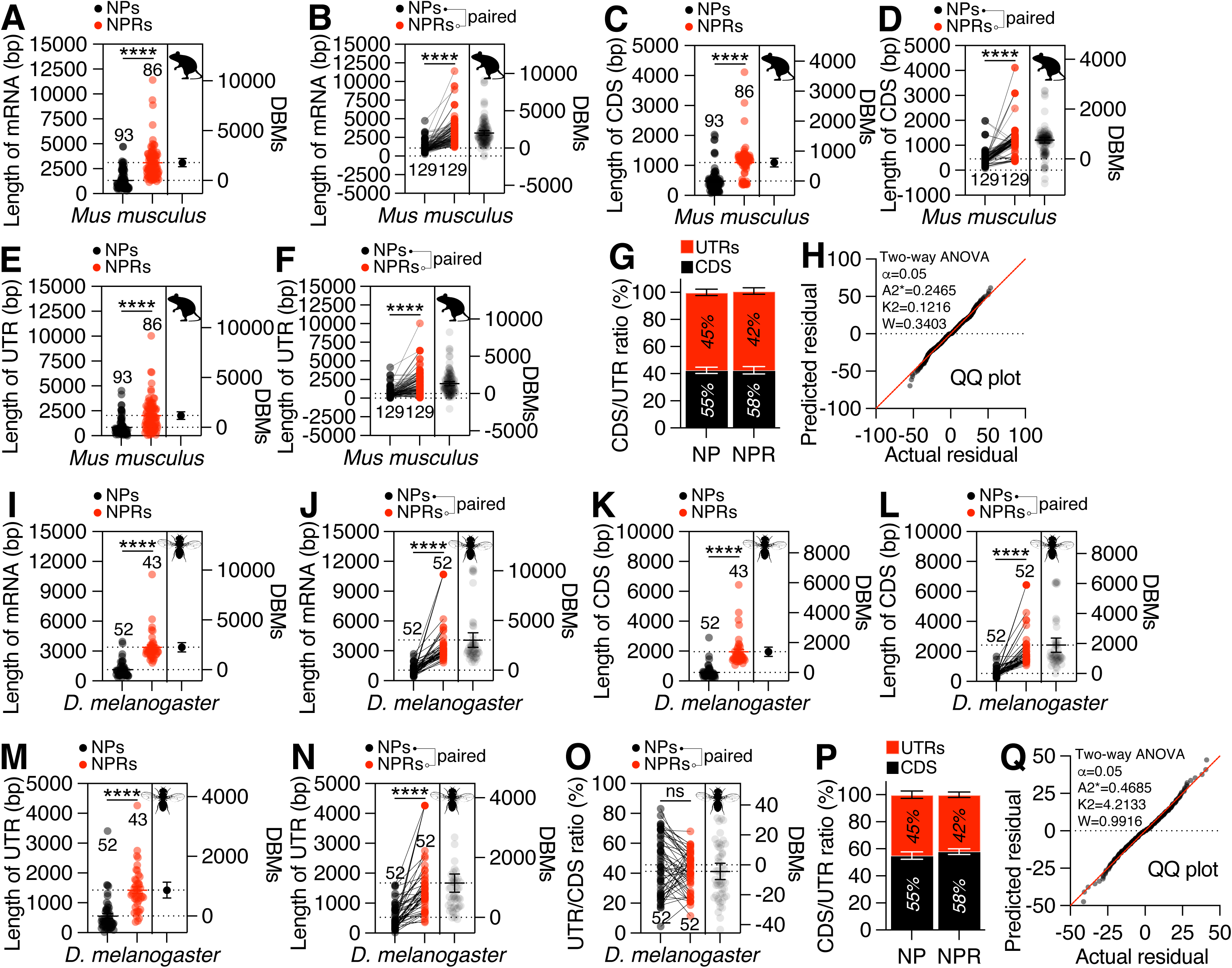
NPR Genes Exhibit More Complex Post-transcriptional Regulatory Landscape than NP Genes across *M. musculus* and *D. melanogaster*. **A-H.** Post-transcriptional Regulatory Landscape in *Mus musculus* displaying total length of mRNA (a-b), CDS (c-d), and UTR (e-f) of NP and NPR genes, along with the DBMs between NP and NPR genes. (g) NP and NPR genes have similar CDS/UTR ratio. (h) QQ plot indicates that the data adhere to a normal distribution. α, Significance level; K2, D’Agostino-Pearson omnibus, A2*, Anderson-Darling; W, Shapiro-Wilk. **I-Q**. *D. melanogaster* Gene Components Landscapes. Post-transcriptional Regulatory Landscape in *D. melanogaster* displaying total length of mRNA (a-b), CDS (c-d), and UTR (e-f) of NP and NPR genes, along with the DBMs between NP and NPR genes. (o-p) CDS/UTR ratio has no significant difference between NP and NPR genes. (q) QQ plot indicates that the data adhere to a normal distribution.

The Coding Sequence (CDS) is the region of mRNA that is translated into a protein. Its length directly correlates with the protein’s size and complexity. The CDS length can also influence mRNA secondary structure and the efficiency of translation, as longer ORFs might affect ribosome traffic and the rate of translation (Narula et al. 2019). The CDS of NPR genes are markedly longer than those of NP genes across both murine and *Drosophila* species, irrespective of whether the genes are unpaired or paired (Fig. 2E–H).

The untranslated regions (UTRs) of messenger RNA (mRNA), found at both the 5’ and 3’ ends of the transcript, are essential modulators of gene expression, including modulation of the transport of mRNAs out of the nucleus and of translation efficiency (Velden and Thomas 1999), subcellular localization (Jansen 2001) and stability (Bashirullah et al. 2001). Strikingly, the lengths of the UTRs of NPR genes markedly exceed those of NP genes, irrespective of whether they are examined in unpaired or paired contexts (Fig. 2I-L). Given the pivotal role of both the 5’ and 3’ UTRs in shaping the post-transcriptional regulatory landscape of mRNA (Mayr 2019; Wieder et al. 2024), these findings imply that NPR genes may represent more intricate targets for cellular post-transcriptional regulatory mechanisms.

The ratio of UTR length to CDS length serves as an indicator of the relative abundance of regulatory elements in comparison to the coding sequence within an mRNA molecule. An elevated UTR-to-CDS ratio is often correlated with an enhanced propensity for post-transcriptional regulation, as it suggests a higher density of regulatory elements that can modulate mRNA processing, stability, and translation efficiency. Empirical evidence has demonstrated that genes characterized by a high 3’UTR-to-CDS ratio are predominantly associated with GO terms that are specific to cell-type functions, reflecting a role in specialized cellular processes (Ji et al. 2021). In the context of the current study, the UTR-to-CDS ratios of NP and NPR genes exhibit a comparable and notably high magnitude in both murine and *Drosophila* species (Fig. 2M-Q). This observation implies that both NP and NPR genes are likely to be implicated in cell-type specific functions, underscoring their potential involvement in specialized regulatory networks within specific cellular contexts.

### Analyzing Neuropeptide and Their Receptors associated with Transcription Factor Network from *Drosophila* Single Cell RNA Sequencing Data

Utilizing the rich genetic repertoire of *Drosophila melanogaster*, a cornerstone model organism in neuroscience, we have harnessed over a century of accumulated datasets to delve into the intricate regulatory mechanisms governing NP and NPR signaling (Bellen et al. 2010; Venken et al. 2011; Attrill et al. 2016; Gramates et al. 2022; Jenkins et al. 2022; Li et al. 2022a; Öztürk-Çolak, Marygold, Antonazzo, Attrill, Goutte-Gattat, Jenkins, Matthews, Millburn, Santos, Tabone, Perrimon, et al. 2024). These signaling pathways are pivotal for orchestrating a spectrum of neurobiological behaviors and physiological responses in the fruit fly (Nässel and Winther 2010; Caers et al. 2012b; Elphick et al. 2018; Nässel and Zandawala 2022). By extracting foundational datasets from the Fly Cell Atlas, encompassing single-nucleus RNA sequencing (snRNA-seq) across the entire organism and specific tissues, we embarked on a comprehensive bioinformatic analysis (Li et al. 2022a; Lu et al. 2023a).

This analysis targeted the intricate interplay between transcription factors (TFs) and the NP-NPR axis, aiming to elucidate the molecular underpinnings of their regulatory networks. Our approach involved the initial aggregation of snRNA-seq data, followed by rigorous computational processing to discern the distinct associations that TFs forge with NPs and NPRs, thereby enhancing our understanding of their role in modulating neurobiological functions.

In our study, we explored the complex relationships between different components of the nervous system using a cutting-edge genetic technique known as single-cell RNA sequencing (snRNA-seq). This method allowed us to examine the activity of genes within individual cells, providing a detailed view of how they interact and contribute to various functions.

We collected data from 17 different tissues of the *Drosophila melanogaster*. The data was sourced from the fly cell atlas portal, which offers comprehensive genetic information in a user-friendly format (Li et al. 2022a; Lu et al. 2023a). Our team employed Python, a popular programming language, to process and organize this vast amount of data. By using specialized software packages, we were able to extract and catalog details about various tissues, specific genes, and individual cells, storing this information in a structured database. Our dataset encompassed a remarkable 16,373 distinct genes across over 500,000 cells. The information was initially in a complex matrix format, with cells and genes arranged in rows and columns, much like the layout of a spreadsheet. This included not only the identity of the genes but also how actively they were being used, or “expressed”, in each cell.

To make sense of this intricate dataset, we used a Python package called Scanpy, which is a scalable toolkit for analyzing large-scale single-cell gene expression data (Wolf et al. 2018a). This tool helped us to systematically organize the data into our database, categorizing it by tissue, cell, and gene type. This meticulous organization was crucial for our analysis, as it allowed us to efficiently identify patterns and relationships among different genes and cells. One of our key goals was to understand how groups of genes work together, a phenomenon known as co-expression. By identifying genes that are often active in the same cells, we can infer how they might be functionally linked. This approach led to the discovery of approximately 450 million instances where gene activity could be correlated across our samples.

Our methodical data storage strategy enabled us to trace the activity of specific genes within particular tissues and cells. This not only helped us to understand how genes interact within their cellular environment but also revealed how they might be influenced by other cellular components, such as transcription factors. These are essentially the cell’s managers, controlling which genes are turned on or off (Chowdhury et al. 2020). In summary, our analysis of snRNA-seq data provided a detailed roadmap of gene activity within the fruit fly’s cells. The fly cell atlas platform provides data on 17 tissues, 16,373 genes, and 507,827 cells. There are 255 cell annotations (Li et al. 2022a). On average, each cell exhibits expression of 881 genes. Using these data, we were able to identify relationships between tissues and cells, as well as between cell types and gene expression. From these relationships, we could identify NPs and NPRs related to gene expression influenced by TFs. By applying deep learning to these association data, we developed a model to predict neuropeptide and neuropeptide receptor genes that are affected by specific transcription factors.

### The Spatial Expression Profile of *Drosophila* NP and NPR Genes Across Tissues

NPs and NPRs have long been recognized as pivotal players in the regulation of behavior and physiology, with their diverse array of functions conserved across the evolutionary spectrum. As detailed in the foundational work by Nässel and colleagues, these molecules orchestrate a wide range of physiological processes and behavioral responses in *Drosophila melanogaster*, serving as critical nodes in the complex network of neuronal signaling (Insel and Young 2000; Hewes and Taghert 2001; Nässel and Winther 2010; Caers et al. 2012b; Elphick et al. 2018; Nässel and Zandawala 2019; Nässel and Zandawala 2022). However, despite the well-documented roles of NPs and NPRs, the underlying mechanisms that govern their expression and activity within the cellular and molecular milieu remain largely enigmatic.

In particular, the intricate interplay between NPs, NPRs, and the TFs that regulate their gene expression has not been extensively explored. This regulatory network is hypothesized to be a key determinant of the dynamic and context-dependent nature of NP-NPR signaling, with profound implications for understanding the nuanced control of behavior and physiology. In this study, we endeavor to chart the landscape of NP-NPR-TF interactions for the first time, shedding light on how the gene expression context modulates the expression dynamics of NPs and NPRs, and by extension, their impact on various behaviors and physiological states. By doing so, we aim to unravel the molecular underpinnings of NP-NPR signaling and its role in mediating adaptive responses to diverse environmental contexts.

To investigate this interaction, we analyzed NPR gene expression across different tissues in fruit flies using single-cell RNA sequencing. Notably, the head region exhibited the highest levels of NPR activity, indicating its pivotal role in NP signaling (Fig. 3A). The correlation between gene expression levels and the number of expressing cells was modest (R = 0.6572; Fig. 3B), suggesting additional regulatory factors at play. When categorized by functional tissue annotation, a strong linear relationship emerged between NPR gene expression and cell counts (Fig. 3C-D). This implies that NPR gene activity and the number of cells expressing NPRs are more closely linked when we consider the functional roles of the tissues, rather than just their physical location. These findings provide a clearer picture of NPR dynamics, underscoring the head as a key area for NP-NPR action and hinting at complex regulatory mechanisms that govern NPR expression.

**Figure 3.**
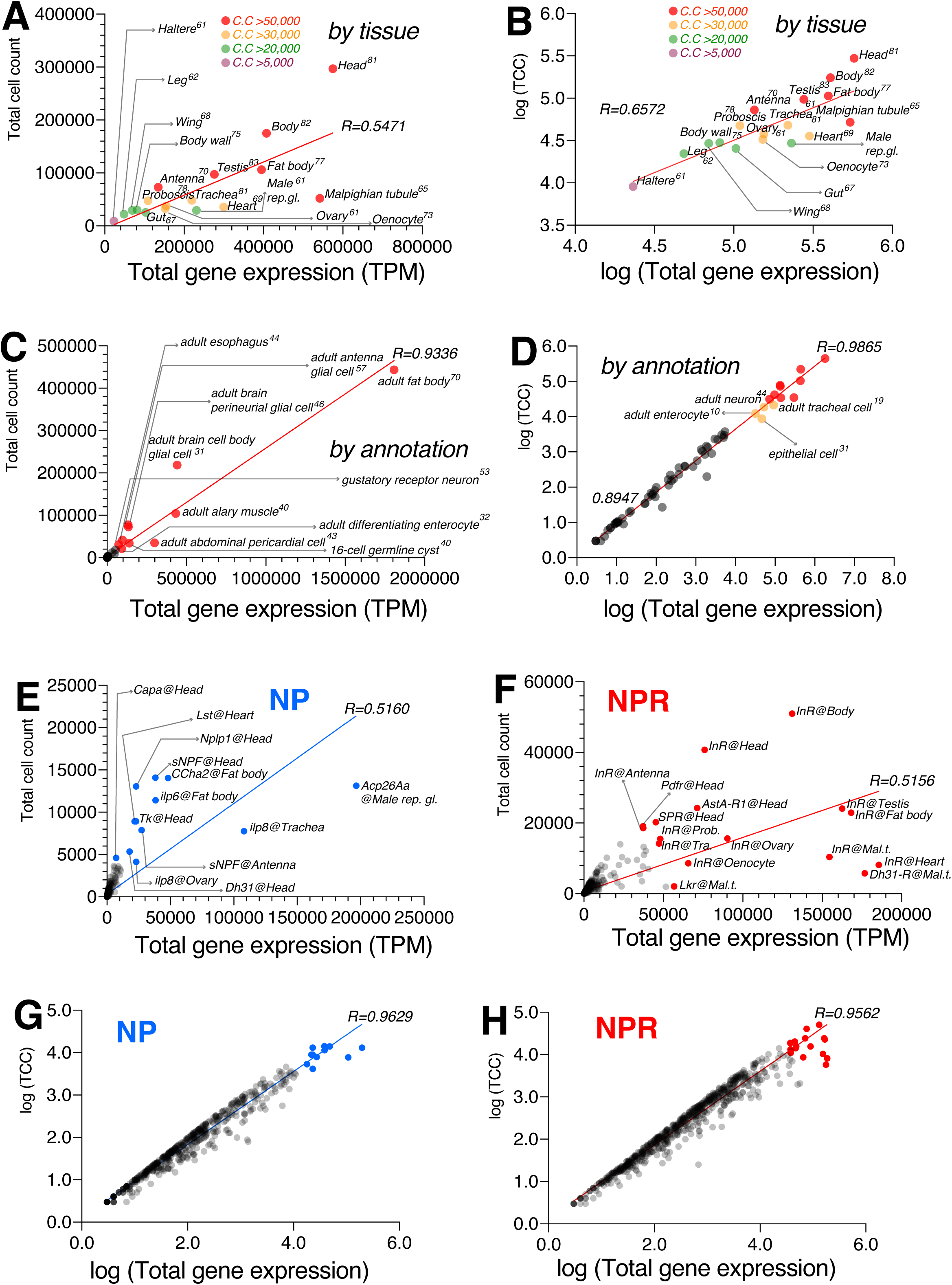
Spatial Expression Profiles and Correlation Analysis of NP and NPR Across *D. melanogaster* Tissues. **A-B**. Scatter Plot of Total Gene Expression (TPM) and Total Cell Count (TCC) by Tissue in *D. melanogaster*, showing that head exhibit the highest expression level compared to other tissues. Tissues are color-coded by cell count (C. C) threshold. Correlation coefficient R indicating its modest correlation. **C-D**. Strong linear relationship of TPM and TCC by annotations in *D. melanogaster*. **E-F**. Scatter plots of TPM versus TCC, highlighting representative high-expression NP (e) and NPR (f) genes, underscoring the role of NP-NPR signaling in essential functions. Notably, Acp26A shows high expression in the male reproductive gland, while sNPF, Nplp1, Tk, Dh31, and Capa are predominantly expressed in the head. InR is the most abundantly expressed receptor, particularly in the body and heart, emphasizing its role in systemic signaling. **G-H**. Strong linear relationship between TPM and TCC for both NP and NPR genes across different tissues in *D. melanogaster*.

Next, we examined how individual NPs and NPRs are expressed across various tissues in fruit flies. We found that a specific neuropeptide, Acp26A, was particularly abundant in the male reproductive gland, highlighting its potential role in reproduction. Other NPs, such as sNPF, Nplp1, Tk, Dh31, and Capa, were prominently expressed in the head, while ilp6 (Bai et al. 2012) and CCHa2 (Li et al. 2013; Ren et al. 2015; Sano et al. 2015) were notably active in the fat body—known for energy metabolism. Notably, sNPF’s expression in the antenna, which is involved in olfaction, aligns with previous studies (Nässel et al. 2008) and validates our approach (Fig. 3E).

Conversely, the expression of NPRs showed a different pattern, with InR—known to bind multiple insulin-like peptides (ilp2, 3, 5, and 6)—being the most prevalent (Graze et al. 2018). Its high expression in the body and heart further underscores the importance of NP-NPR signaling in vital functions (Fig. 3F). Our data revealed a strong positive correlation between gene expression levels (TPM) and the number of cells expressing these genes (TCC) for both NPs and NPRs, indicating that tissues with high gene expression also contained more cells expressing these genes (Fig. 3G-H).

These insights suggest that the abundance of certain NPs and NPRs is closely linked to critical functions like reproduction and survival, with the brain-to-gut-to-fat body axis emerging as a key pathway for NP-NPR signaling in these processes (Fig. S3) (Nässel and Winther 2010; Wielendaele et al. 2013; S.M. Kim et al. 2016; Nässel and Zandawala 2019; Semaniuk et al. 2021; Nässel and Zandawala 2022).

### The Landscape of *Drosophila* NP-NPR-TF Network

To elucidate the regulatory network involving TFs, NPs, and NPRs in modulating context-dependent behaviors, we focused on 10 NPs and 13 NPRs implicated in interval timing behaviors (Table 2). These selections align with our objective to investigate the modulation of complex behaviors by a limited set of NP-NPR pairs, as evidenced in prior studies (Kim et al. 2013; Wong et al. 2019; W.J. Kim et al. 2024a; T. Zhang et al. 2024a). Leveraging the AUCell algorithm from the fly cell atlas, we harnessed the spatial resolution of TF data to visualize and analyze the NP-TF and NPR-TF interactions, shedding light on the transcriptional mechanisms that sculpt the NP-NPR network architecture (Aibar et al. 2017a; Sande et al. 2020).

In our analysis of the transcriptional landscape, we observed a distinct pattern of association between NPs, NPRs, and their corresponding TFs (Fig. 4A). A visualization of 10 NPs and 13 NPRs along with their TF network revealed that cells expressing NPRs exhibit a significantly higher expression of TFs (Fig. 4B). Comparative analysis of network nodes associated with either NPs or NPRs demonstrated a higher connectivity for TFs linked to NPRs (Fig. 4C), indicating a more complex and diverse transcriptional regulation in NPR-expressing cells.

**Fig. 4.**
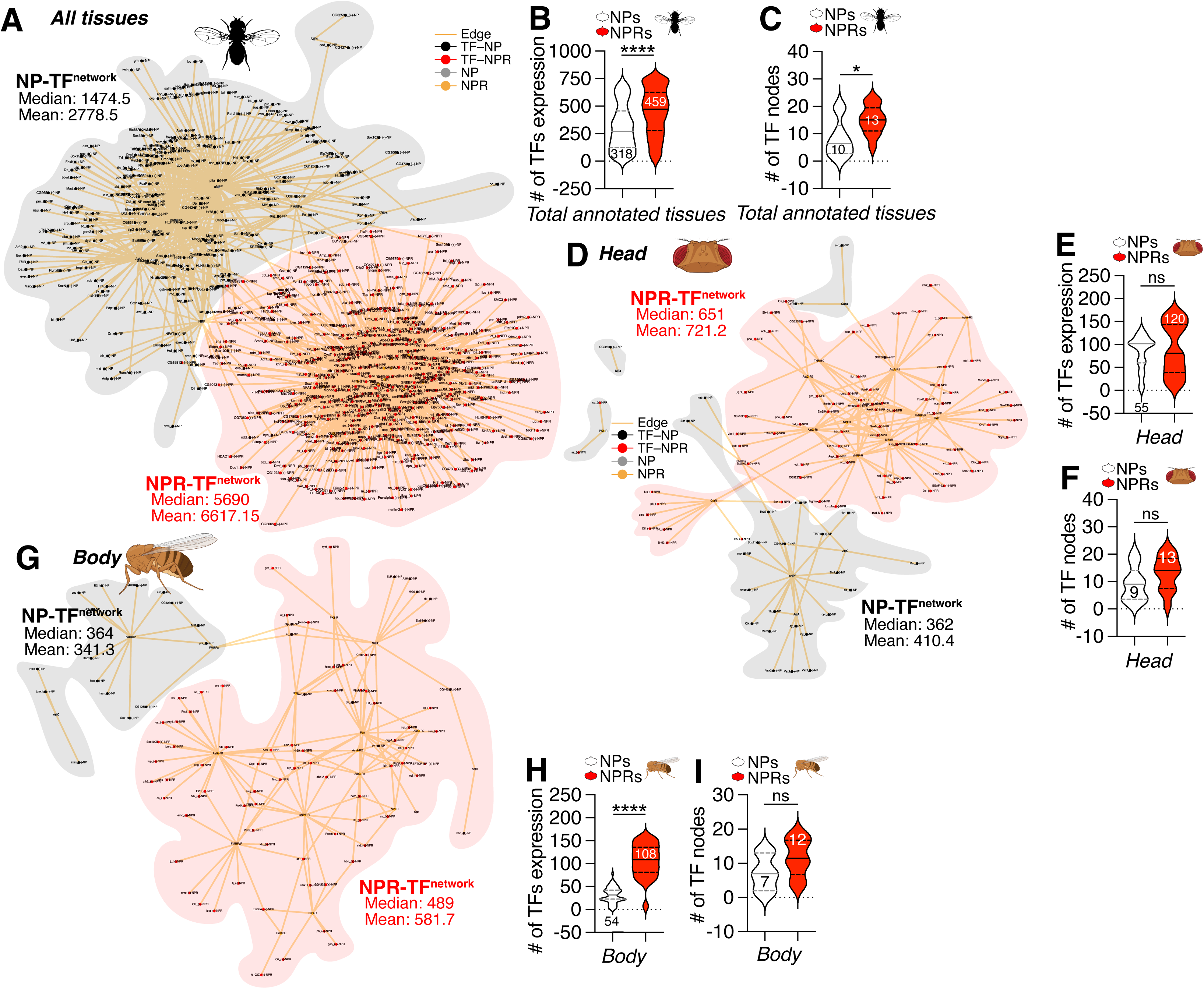
The relationship between TFs that regulate the transcription of 10 NPs and TFs that regulate the transcription of 13 NPRs. **A:** The overall tissue-independent network of TFs that regulate NP transcription and TFs that regulate NPR transcription across 17 tissues. The brown connecting edges represent the relationship between NPs or NPRs and TFs involved in transcription regulation. Black dots represent TFs that regulate the transcription of NPs. Red dots represent TFs that regulate the transcription of NPRs. Gray dots represent NPs that are regulated by TFs. Brown dots represent NPRs that are regulated by TFs. The gray mesh represents the network between NPs and TFs, while the brown mesh represents the network between NPRs and TFs. The median and mean values were calculated from the total number of individually expressed instances of NPs and NPRs. The data was constructed by using the Motif Regulon information from each of the 17 tissue cells to identify and connect NPs, their transcriptional regulators (TFs), NPRs, and their transcriptional regulators (TFs) within the same cell. For detailed information, please refer to the “NP, NPR, TF Network” section in the **MATERIALS AND METHODS**. **B**: Number of TFs regulating NP and NPR. **C**: Number of NPs and NPRs regulated by TFs. **D**: Network of TFs that regulate NPs and NPRs in the head (The details are the same as in A.). **E**: Number of TFs regulating NP and NPR in the head. **F**: Number of NPs and NPRs regulated by TFs in the head. **G**: Network of TFs that regulate NPs and NPRs in the body (The details are the same as in A.). **H**: Number of TFs regulating NP and NPR in the body. **I**: Number of NPs and NPRs regulated by TFs in the body. The graphical representation of the data (B-C, E-F, H-I) was executed using a Ranks plot, which is a non-parametric method employed after the Mann-Whitney U test. This approach was selected due to the non-normal distribution of the TF expression and TF node values. Consistent with this methodology, all subsequent analyses and visualizations throughout the manuscript will adhere to this statistical framework. For an exhaustive delineation of the statistical procedures employed, readers are directed to the **MATERIALS AND METHODS** section of this manuscript.

When we narrowed our focus to the head tissue, the pronounced dominance of NPRs in TF association was no longer evident (Fig. 4D). Although TF expression in NPR-expressing cells remained higher than in NP-expressing cells (Fig. 4E), and the node count was consistent (Fig. 4F), these differences were not statistically significant, suggesting a comparable level of transcriptional regulation for NPs and NPRs in the head. In stark contrast, the body’s NPR-TF network exhibited a significantly higher complexity than the NP-TF network (Fig. 4G). Cells expressing NPRs displayed an exceptionally higher TFs expression (Fig. 4H), yet the disparity in network nodes did not reach statistical significance (Fig. 4I), pointing to a more intricate interplay between TFs and NPRs in the body’s transcriptional landscape.

Our findings reignite the long-standing query regarding the multifaceted roles of NPs, which often moonlight as hormones. Unlike NTs that operate chiefly within the confines of synapses, typically between neurons and, in some cases, muscle or glial cells, NPs exert their influence at a distance by engaging with NPRs that are expressed not only in the brain but also in peripheral tissues throughout the body. In humans, NPs can modulate the efficacy of synaptic transmission by either potentiating or inhibiting the activity of co-released NTs in the brain, and in the periphery, they can behave akin to peptide hormones, impacting a broad array of physiological functions (Russo 2017).

Although hormones are distinct from NPs (Zup et al. 2022), many NPs were first identified as hormones from the pituitary gland or gastrointestinal tract (Dawson 1999; Siegel 1999). This dual functionality may stem from the fact that NPs are the most structurally and functionally diverse group of chemical messengers. They are produced by neurons in the nervous system and by neurosecretory cells, earning them the functions of neuropeptides or peptide hormones (PHs), respectively. Moreover, many peptides are synthesized by endocrine cells or other cell types in various locations, with a single peptide capable of being expressed by any of these cell types within an organism. The omnipresence of NPs and PHs in the nervous and endocrine systems of all metazoans underscores their critical roles.

Given that NPRs are the gateways for both NPs and PHs, the complexity of the NPR-TF network in the body over the head is not surprising. In the fruit fly *Drosophila melanogaster*, the neuropeptide AstA and NPF are both expressed in the brain and gut, where they serve as neuropeptides and hormones. These peptides exert their effects through the same NPRs, which are widely expressed (Malita et al. 2022; Li et al. 2023; Gao et al. 2024). This complexity may reflect the diverse physiological demands across bodily systems compared to the head. Recent reports have indicated that different NPRs can distinctly regulate behaviors such as movement in *C. elegans*, with varying expression patterns in the head and body (Ramachandran et al. 2021). This underscores the rationale for our observation of a more intricate NPR-TF network in peripheral tissues, likely due to the broader range of physiological roles that NPs and their receptors play outside the central nervous system.

Upon the exclusion of TFs that are co-expressed in both NPs and NPRs, the network patterns of NP-TF and NPR-TF became more distinctly discernible (Fig. S4A). The removal of TFs common to NP- or NPR-expressing cells revealed a persistent bias toward NPRs, indicating that NPR-expressing cells engage in a more intricate network with TFs (Fig. S4B). It is a well-established fact that the cascade of TFs is a key feature in controlling developmental processes in *Drosophila melanogaster* (Bolouri and Davidson 2003). The TF cascade is also essential for generating diversity and plasticity in enteroendocrine cells (Guo et al. 2022). Notably, TF cascades are critical control mechanisms for neuronal diversity (Sousa and Flames 2022), synaptic plasticity (Engelmann and Haenold 2016), and long-term memory formation (Alberini 2009). This raises the possibility that TF cascades may also be responsible for modulating neuronal and behavioral plasticity in interval timing. As anticipated, the prevalence of TFs regulated by other TFs, inferred as part of a TF cascade, was significantly higher in NPR-expressing cells compared to NP-expressing cells, both in terms of network motifs and tissue context (Fig. S4C-D). These findings strongly suggest that TFs exclusively expressed in NPR-expressing cells are highly regulated by TF cascades and may thus control the behavioral plasticity necessary for NP modulation of multiple context-dependent behaviors.

When considering TFs that are not shared between NP- and NPR-expressing cells, both head and body TF networks exhibited a pronounced bias toward NPR-expressing cells, with a significantly higher number of TFs present in these cells, particularly in the body (Fig. S4E-H), as previously indicated in Fig. 4D-I. Consequently, these data further support the notion that the TF network is more complex in NPR-expressing cells compared to NP-expressing cells, and in the body compared to the head.

### Revealing Tissue-Specific Bias in Gene Regulatory Networks: The Dominant Role of NPR-TF

We conducted a comprehensive assessment of regulatory networks in various tissues, focusing initially on internal organs. It is well established that transcriptional regulation plays a pivotal role in maintaining metabolism and responding to infections within these organs (Clayton 2002; Desvergne et al. 2006; Hirota and Fukamizu 2010; Mammoto et al. 2012; Mandal et al. 2018; Scholtes and Giguère 2022). Across all internal organs examined, including the fat body, gut, heart, Malpighian tubules, and oenocyte, we found that cells expressing NPR exhibited a higher number of TFs compared to those expressing NP. With the exception of the trachea, the difference in TF expression levels was statistically significant, indicating a pronounced regulatory control favoring NPR-expressing cells (Fig. S5).

We also explored the transcriptional control in sensory organs, which are known to be highly regulated at the transcriptional level (Weinberger et al. 2017). Similar to internal organs, most sensory organs showed a marked increase in TFs associated with NPR-expressing cells. However, a notable exception was observed in the antenna, where NP-expressing cells displayed higher TF expression levels (Fig. S6A-C). This unique pattern may be attributed to the rich expression of short neuropeptide F (sNPF) in the antennal lobe (AL) of adult flies (Carlsson et al. 2010), which is crucial for olfactory sensory cell function (Nässel et al. 2008; Carlsson et al. 2013). Our data also suggest that sNPF is a dominant factor in the TF network of NP-expressing cells in this tissue (Fig. S6A).

In the context of sexually dimorphic organs, we discovered an interesting bias in the male reproductive gland, where NP-expressing cells showed significantly higher TF expression. This could be linked to the influence of accessory gland proteins (Acps), which are known to affect female physiology and behavior (Wolfner 2002; Chapman and Davies 2004; Ram and Wolfner 2007). Notably, Acp26a was highly expressed in the male reproductive gland among the NP genes (Fig. 3E). Despite this, the overall TF network and the number of TF nodes were more abundant in NPR-expressing cells, suggesting a complex regulatory interplay. In contrast, the female ovary and male testis followed the pattern observed in other tissues, with NPR-expressing cells being more tightly regulated by the TF network. This consistency across genders indicates that the bias towards an NP-biased TF network in the antenna and male reproductive organ is an exception rather than the rule (Fig. S7).

Upon removing TFs common to both NP- and NPR-expressing cells, we identified only six tissues where NP-expressing cells retained unique TFs: the gut, fat body, body wall, heart, oenocyte, and wing. In each of these tissues, the number and expression levels of TFs were higher in NPR-expressing cells (Fig. S4I). This finding implies that NP-expressing cells largely rely on a shared set of TFs that are also utilized by NPR-expressing cells across most tissues, with a limited number of TFs being exclusively expressed in NP-expressing cells.

### The neuronal populations that express genes encoding NPs and their corresponding NPRs share a remarkable degree of biochemical homogeneity

The neuropeptidergic and neurotransmitter systems represent two pivotal modalities of cellular communication within the nervous system, each endowed with unique properties and roles (Nässel and Winther 2010; Schoofs et al. 2016; Nässel 2018; Nässel and Zandawala 2022). Neuropeptides, a diverse and elongated class of signaling molecules, are distinguished from conventional neurotransmitters by their capacity to modulate physiological functions over extended timeframes. Their multifaceted influence on behavior and metabolism stems from their widespread action potential across the organism. Conversely, neurotransmitters, which are low-molecular-weight compounds, are emitted from neurons to facilitate rapid signal transmission across synaptic clefts to neighboring neurons or target cells. They are integral to swift, short-lived interactions within the nervous system and are essential for functions such as muscle activation, sensory perception, and cognitive processing. The neurotransmitter system is renowned for its expeditious and targeted signal conveyance across brief intervals (Bhat et al. 2021).

The neurochemical signaling network in *Drosophila melanogaster* exhibits parallels with that of other species, including *Homo sapiens*, and is integral to the proper functioning of the nervous system (Martin and Krantz 2014). Within this model organism, a diverse array of NTs is present, including dopamine (DA), gamma-aminobutyric acid (GABA), glutamate (Glu), serotonin (5-HT), acetylcholine (ACh), octopamine (OA), tyramine (TA), and histamine, each contributing significantly to the modulation of physiological processes and behavioral patterns (Shin et al. 2018). Although comprehensive studies on NTs in *Drosophila* are limited, it is well-documented that many neurons in this species are capable of synthesizing more than one type of NT (Croset et al. 2018; Nässel 2018). Neurotransmitters typically interact with multiple receptor subtypes NTRs, eliciting a variety of cellular responses in the target cells (Kondo et al. 2020). Given the analogous nature of the NT-NTR and NP-NPR systems in terms of their regulatory influence on behavior and physiology through neural signaling, we have endeavored to examine the NT-NTR system within a context analogous to our previous investigations of the NP-NPR system.

Utilizing the recently published adult *Drosophila* RNA sequencing dataset, we conducted an analysis to identify cells expressing key enzymes involved in the synthesis of various NTs (Li et al. 2022a). Initially, we surveyed the expression of NT-synthesizing enzymes across 255 annotated cell populations. In cell populations that express NPs but not their NPRs, we observed that cells expressing choline acetyltransferase (ChAT) were the most prevalent, with ChAT expression levels also being relatively high on average (Fig. 5A-B). When assessing the total cell count, overall gene expression, and the number of tissues expressing NT-synthesizing enzymes, histidine decarboxylase (Hdc), which is crucial for the synthesis of histamine, demonstrated the highest abundance (Fig. 5C-E). In cells expressing NPRs, there was a greater number of cells expressing NT-synthesizing enzymes, along with higher gene expression levels for these enzymes (Fig. 5F-G), likely reflecting the fact that the number of cells expressing NPRs exceeds those expressing NPs. Cells expressing NPRs also exhibited a robust Hdc expression profile (Fig. 5H-J). Despite variations in the absolute levels of NT-synthesizing enzymes, the overall pattern of enzyme expression was similar between cells expressing NPs and those expressing NPRs (Fig. 5A-J), indicating that the synthesis of NTs in cells expressing either NPs or NPRs is not significantly different. Collectively, these findings suggest that the neurochemical landscape of neurotransmitter expression within cell populations expressing NPs or NPRs is strikingly similar.

**Figure 5.**
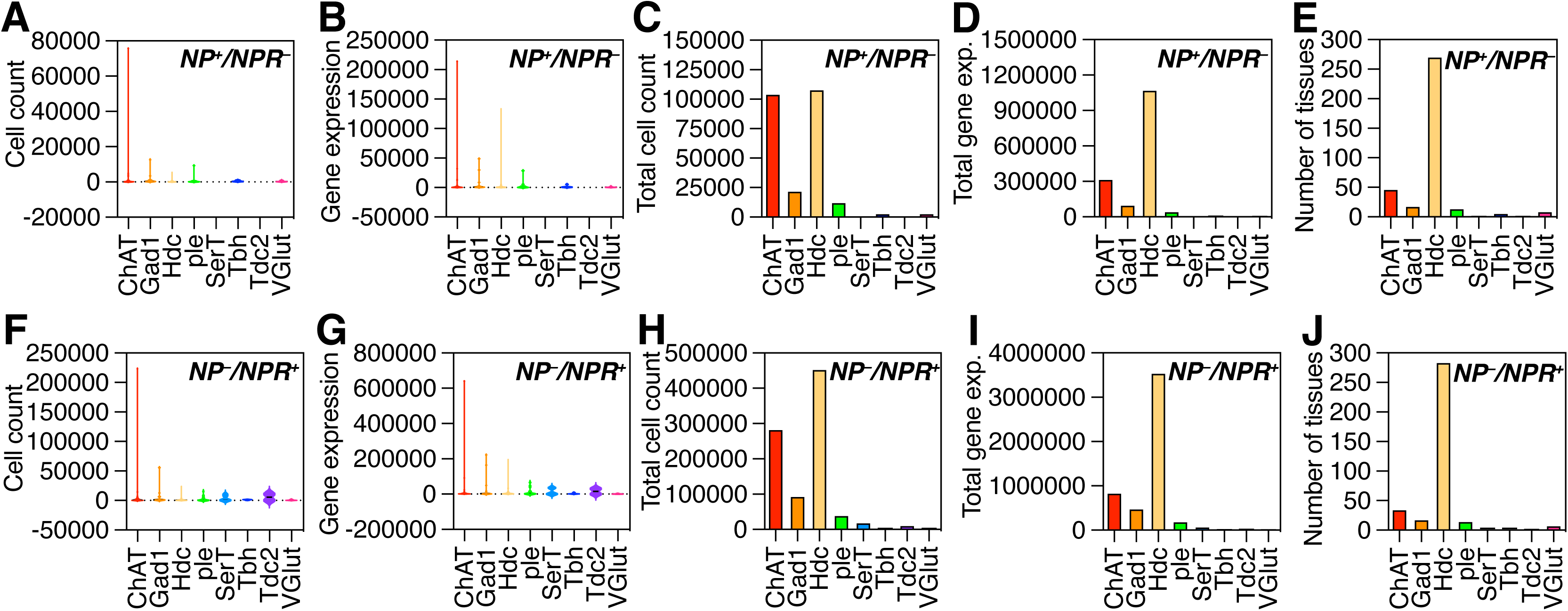
Similar Neurochemical landscape of neurotransmitter (NT) expression between cells expressing NPs and cells expressing their corresponding NPRs in *D. melanogaster*. **A-E.** Neurochemical landscape of NT expression in cells expressing NPs but lacking NPRs. (a-b) Cell count and gene expression levels of NT-synthesizing enzyme genes in cells expressing NPs but not the corresponding NPRs, with each dot representing a distinct cell type. (c-e) Total cell count, aggregate gene expression, and number of positive tissues in NT-synthesizing enzyme-positive NP cells. **F-J. Neurochemical landscape of NT expression in cells expressing NPRs but lacking NPs.** (a-b) Cell count and gene expression levels of NT-synthesizing enzyme genes in cells expressing NPRs but not the corresponding NRs. (c-e) Total cell count, aggregate gene expression, and number of positive tissues in NT-synthesizing enzyme-positive NPR cells.

Given our observation that cells expressing NPRs are subject to biased transcriptional regulation, we sought to explore the TF-associated network between cells expressing NTs and those expressing their NTRs. The identity of NTs within a neuron is a fundamental characteristic that must be stringently controlled to ensure the development of a functional nervous system. It is well established that the tissue content of NTs fluctuates dynamically with the life stages of *Drosophila.* For instance, the levels of TA and OA undergo significant changes during the development of the fruit fly, with elevated TA but reduced OA levels observed in pupae. In adult *Drosophila*, DA tissue content is notably higher in females compared to males (Denno et al. 2015). It has been extensively documented that the expression of NTs in neurons is tightly regulated by specific TFs during development (Estacio-Gómez et al. 2020). Notably, studies have identified and emphasized the differential expression of TFs and non-coding RNAs among various types of NT neurons. In particular, TFs enriched in cholinergic neurons are expressed in largely non-overlapping populations within the adult brain, suggesting the absence of combinatorial regulation of NT fate in this context (Estacio-Gómez et al. 2020).

In our analysis of the TF networks associated with NT-expressing and NTR-expressing cells, we observed no significant difference in the total number of TFs between these two cell types (Fig. S9A). However, the overall expression of TFs was markedly higher in NTR-expressing cells (Fig. S9B). The number of nodes within the TF network was also higher in NTR-expressing cells, although this difference did not reach statistical significance compared to NT-expressing cells (Fig. S9C). A comparative analysis of representative tissues, such as the head and body, revealed substantial differences in the landscape of TF networks when contrasted with the NP-NPR-TF network.

In head tissue, the number of TFs expressed in both NT- and NTR-expressing cells was similar, but the number of cells expressing these TFs was significantly greater in NT-expressing cells than in NTR-expressing cells (Fig. S9D-E). The number of nodes within the TF network was also significantly higher in NT-expressing neurons compared to NTR-expressing cells (Fig. S9F). When compared to the NP-NPR-TF network analysis in the head (Fig. 4D-F), these findings suggest that NT-expressing neurons are subject to more stringent transcriptional control by the TF network. A similar phenomenon was observed in body tissue, where there were no significant differences in the number of TF-expressing cells or TF network nodes between NT- and NTR-expressing cells (Fig. S9G-I), indicative of a highly biased TF regulatory network in NPR-expressing cells (Fig. 4G-I). Consequently, in the context of TF networks for NT- and NTR-expressing cells, our data indicate that NT-expressing cells are more tightly regulated by a dense network of TFs.

This observation prompts the intriguing query of how the NP-NPR and NT-NTR systems have evolved distinct landscapes for the transcriptional control of their regulatory networks by TFs. It suggests that in the NP-NPR system, cells expressing the receptor are subject to intense TF network regulation, whereas in the NT-NTR system, cells expressing the ligand are under tight TF network control. These divergent TF-dependent regulatory mechanisms of the NT-NTR and NP-NPR systems imply that intricate regulatory networks have co-evolved to fine-tune complex and context-dependent behavioral repertoires through TF-mediated transcriptional regulation.

Gene transcription is the fundamental process through which the genetic information of an organism is deciphered into a series of directives for cellular activities such as division, differentiation, migration, and maturation. As cells operate within their specific environments, transcription also enables mature cells to engage dynamically with their surroundings while steadfastly preserving essential information about historical experiences. Given the significance of this process, it is reasonable to consider that activity-dependent transcription in the vertebrate brain is a key mechanism, with contemporary research encompassing studies on activity-dependent chromatin modifications and the plasticity mechanisms that underpin adaptive behaviors (Yap and Greenberg 2018). Understanding how these neuronal activity-dependent transcriptional mechanisms contribute to our comprehension of the refinement and plasticity of neural circuits for cognitive function is not only crucial for vertebrates but also for invertebrate brains, helping to bridge gaps in our knowledge regarding these processes.

### Transcriptional regulation of NP signaling in aging

Aging is a complex biological process characterized by the gradual decline in physiological function and an increased susceptibility to age-related diseases. At the molecular level, aging is associated with significant alterations in gene expression, which are orchestrated by transcriptional control mechanisms. TFs play a crucial role in these processes, as they bind to specific DNA sequences and regulate the expression of target genes involved in cellular maintenance, stress response, and metabolic regulation (Maity et al. 2022). The transcriptional control of aging is modulated through a dynamic interplay of various TFs and signaling pathways. Among the key factors are those involved in the insulin/insulin-like growth factor 1 signaling (IIS) pathway, the target of rapamycin (TOR) pathway, and the FOXO family of forkhead transcription factors. These factors influence lifespan and healthspan by regulating the expression of genes related to antioxidant defense, DNA repair, autophagy, and mitochondrial function (Roy et al. 2002; Stegeman and Weake 2017; Tia et al. 2018).

Based on our findings that transcriptional control is more biased in NPR-expressing cells, we tried to confirm these TF-biased control can be confirmed in age-related changes in adult *Drosophila*. To identify the TFs that regulate NP and NPR gene expression across different ages, we first used regulon data from the FCA. Several TFs were identified as regulators of both NP and NPR genes, using the Aging Fly Cell Atlas (AFCA) (Lu et al. 2023a) to examine changes in their expression across the head and body. Notably, more TFs regulate NPRs than NPs, with minimal overlap between the two categories (Fig. 6A).

**Figure 6.**
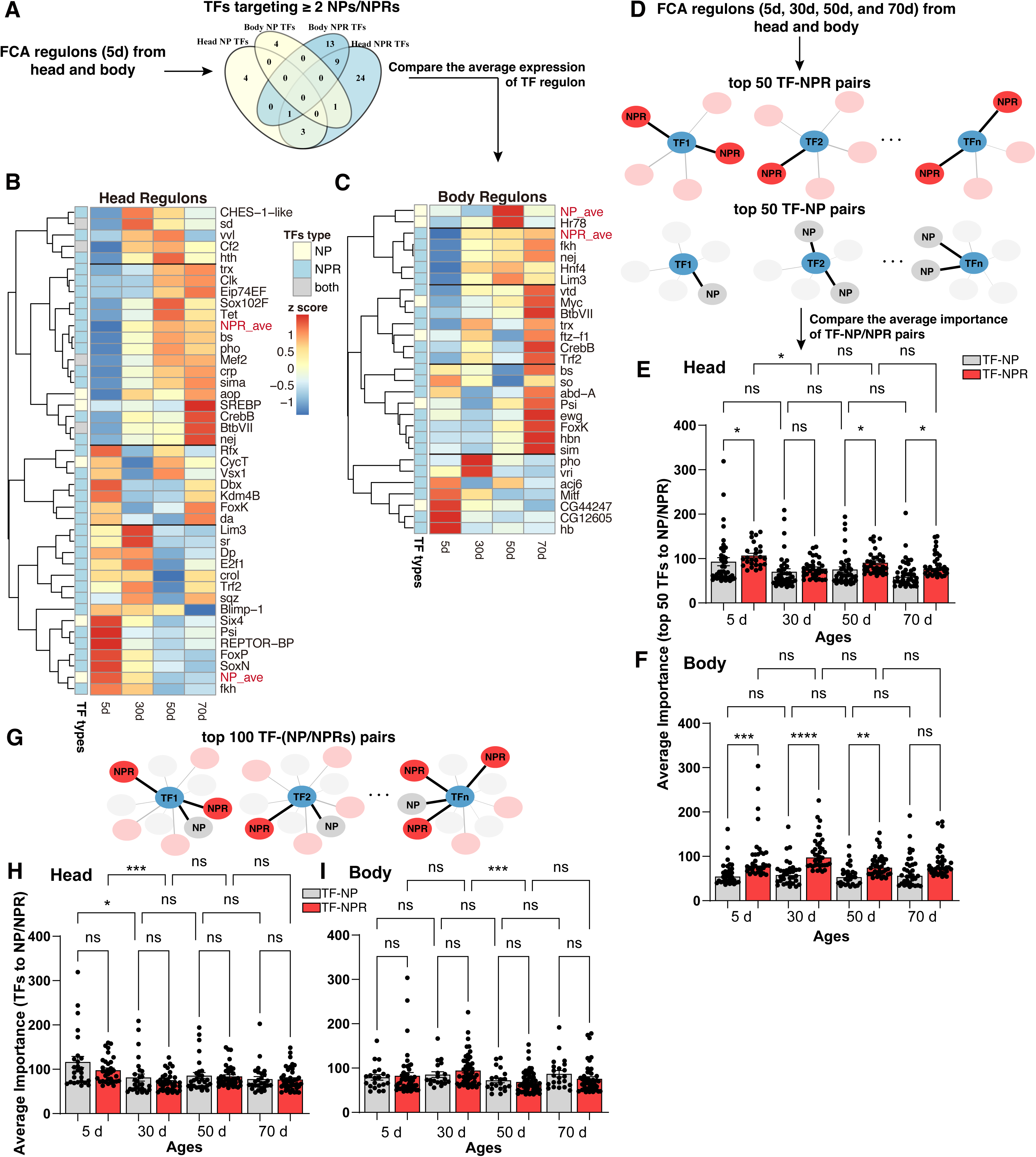
Identification and Analysis of TFs Regulating NP and NPR Genes Across Aging in *Drosophila*. **A-C**. Identification of TFs involved in the regulation of NP and NPR genes across different ages. (a) Schematic overview of the process for identifying TFs that regulate NP and NPR genes using regulon data from the Aging Fly Cell Atlas (AFCA). The Venn diagram illustrates the overlap between TFs targeting NP and NPR genes in the head and body, showing that more TFs are associated with NPR regulation than NP, with minimal overlap between the two. (b-c) Heatmaps of averaged expression levels of NP- and NPR-related regulons across various time points in the head (b) and body (c), revealing distinct age-related patterns in TF regulation for NPs and NPRs. **D-F**. Analysis of the significance of TF-NP and TF-NPR regulatory pairs over time. (d) Approach used to rank and compare the top 50 TF-NP and TF-NPR pairs based on their importance scores at different ages. (e) Bar plots comparing the average importance of top TF-NP and TF-NPR pairs in the head at ages 5, 30, 50, and 70 days, showing that TF-NPR pairs generally have higher importance scores than TF-NP pairs across these ages, especially in earlier stages. (f) Bar plots comparing the average importance of TF-NP and TF-NPR pairs in the body across the same age points, showing a similar trend of higher importance for TF-NPR pairs at 5, 20, and 50 days, with a slight increase in TF-NPR importance at 70 days. **G-I**. Examination of the top 100 TF-NP and TF-NPR pairs. (g) Network diagram representing the top 100 TF-NP and TF-NPR regulatory pairs, with TF-NPR pairs consistently outnumbering TF-NP pairs, suggesting a more complex regulatory network for NPR genes. (h-i) Bar plots showing the average importance of TF-NP and TF-NPR pairs across different ages in the head (h) and body (i), with results indicating that although NPR genes are regulated by more TFs, the overall importance scores for these pairs do not significantly differ between NP and NPR networks as age progresses.

At the mRNA level, NPR expression increases with age, while NP expression peaks at 5 days in the head and declines at 30, 50, and 70 days. In the body, NP expression is highest at 50 days (Fig. 6B-C). Additionally, slightly more regulons are co-expressed with NPRs than NPs as flies age. Overall, our data depict the expression patterns of NP and NPR genes and their associated regulons, showing distinct trends.

To further explore regulatory differences between NP and NPR genes across different time points, we analyzed TF-gene networks at each age. We ranked TF-NP and TF-NPR pairs by importance and selected the top 50 pairs for comparison (Fig. 6D). In the head, TF-NP pair importance slightly decreases with age, while TF-NPR pair importance shows a stronger downward trend. Notably, at multiple time points, the importance of TF-NPR pairs is significantly higher than that of TF-NP pairs (Fig. 6E). In the body, these differences are more pronounced: TF-NPR pairs exhibit significantly higher importance at 5, 20, and 50 days, while showing a slight increase at 70 days. In contrast, TF-NP pairs maintain a relatively constant importance level, while TF-NPR pairs fluctuate slightly without a clear trend as age progresses (Fig. 6F). These findings suggest that, when focusing on top-ranked (highest importance) TF-gene pairs, the regulatory distinctions between NPR and NP genes become more apparent. This indicates that NPR genes may be more strongly regulated by specific key TFs, whereas NP gene regulation appears more dispersed, lacking similarly high-importance TF-gene pairs.

To gain further insight into the differences between NP and NPR gene regulation, we expanded our analysis by selecting the top 100 TF-NP/NPR pairs (Fig. 6G). Across all ages, both in the head and body, there were consistently more TF-NPR pairs than TF-NP pairs, though the importance scores did not significantly differ between NP and NPR pairs as age increased (Fig. 6H-I). Our results suggest that NPR genes are regulated by more TFs, potentially indicating a more complex regulatory network or a greater degree of cooperative regulation.

In conclusion, NPR genes involve more TFs in their regulation, although the overall regulatory intensity of individual TFs on NPR and NP genes is similar at a general threshold level. However, the differences in the importance of top-ranked pairs suggest that NPR genes are more influenced by key TFs with strong regulatory effects, while NP gene regulation is more spread out and lacks such prominent regulators. This suggests that the regulation of NPR genes may depend more heavily on a few critical TFs, whereas NP gene regulation might be broader but less concentrated on high-importance TFs.

Accordingly, it has become apparent that TF-NPR networks are increasingly significant in the aging process of *Drosophila melanogaster*. To determine whether transcriptional control is also a fundamental component of NP-NPR signaling during early adult development, we utilized scRNA-seq data from 1-day and 3-day old adult flies and compared these with the 5-day old adult NP-NPR-TF networks, which we had previously analyzed (Fig. 4A-C) (Li et al. 2022a). The prevalence of cells expressing NPs peaked at 3 days, whereas the number of cells expressing NPRs continued to rise through to 5 days (Fig. S10A). When assessed at the network level, the complexity of NP-TF and NPR-TF networks was indistinguishable in 1-day old adults (Fig. S10B-C). However, the complexity of the NPR-TF network increased significantly by day 3 (Fig. S10E), and the expression of TFs was elevated in 3-day old adults (Fig. S10F-G). A significant upregulation in the number of TFs was observed between days 3 and 5 (Fig. S10H). Collectively, these findings support the notion that during the early stages of adult development, from day 1 to day 5, which encompasses the period of sexual maturation, NP-NPR signaling is highly regulated by TF networks, particularly in a manner biased towards NPR-TF interactions.

### Diverse transcriptional regulation of NPR expression by distinct NPs-NPRs pairs in modulating context-dependent behaviors

In our study, we observed that the global pattern indicates a stringent transcriptional regulation mediated by TFs over cells expressing NPRs. Nevertheless, the regulation of each NP and its corresponding NPR appears to be associated with distinct TF-based mechanisms. To elucidate the differential regulation of various NP-NPR pairs under TF-based, context-dependent control, we initially focused on the NP-NPR signaling pathway that has been most extensively studied. The insulin signaling pathway is recognized as a critical regulatory hub that sustains metabolic equilibrium and influences a spectrum of health and disease outcomes, including the process of aging. The insulin receptor (IR) initiates a cascade of events that are essential for glucose uptake, glycogen synthesis, lipogenesis, and protein synthesis, as well as the inhibition of gluconeogenesis and lipolysis. Disruptions in insulin signaling can lead to insulin resistance, a condition characterized by a reduced response to insulin, which is central to the pathophysiology of type 2 diabetes (T2D), obesity, cardiovascular disease, and non-alcoholic fatty liver disease (NAFLD) (Desvergne et al. 2006; Mounier and Posner 2006; Hirota and Fukamizu 2010; Scholtes and Giguère 2022).

Insulin signaling plays a crucial role in the regulation of various physiological processes in *Drosophila melanogaster*, including the control of metabolism, growth, and lifespan. The insulin signaling pathway in *Drosophila* involves the insulin receptor (InR), which when activated, phosphorylates and activates the PI3K/AKT pathway. This pathway further affects the activity of the FOXO transcription factor, leading to its exclusion from the cell nucleus, thereby regulating the expression of downstream genes (Nässel et al. 2013; Nässel et al. 2015; Chowański 2021; Semaniuk et al. 2021; Yoon et al. 2022).

To determine the susceptibility of various cell populations to transcriptional regulation, we conducted a comparative analysis of TFs in ilp-expressing cells and InR-expressing cells. The comparative analysis revealed a significantly higher prevalence of TF-expressing cells among InR-positive cells in both the head and body regions (Fig. S11A). Further examination of the TF landscape associated with ilp- or InR-positive cells demonstrated a more intricate network interplay, particularly within the InR-TF cellular networks, along with a markedly greater expression of TFs in InR-expressing cell populations in both head and body (Fig. S11B-E). These findings suggest that cells expressing InR are more likely to be under transcriptional control.

Subsequently, we deeply examined the association of TFs with each NP and NPR pair. Through this comprehensive analysis, we discovered that the majority of NP-NPR pairs exhibit a pattern akin to that of ilps and the InR, indicating that cells expressing NPRs are more stringently governed by TF-mediated transcriptional regulation within TF-NPR networks (Fig. S11-S14). Nonetheless, we identified several outliers, suggesting that cells expressing NPs may be more vulnerable to TF-associated transcriptional control. For instance, when analyzing the AstA-AstA-R1 pair, we observed a canonical NPR-biased expression of TFs (Fig. S11F-G). In contrast, cells expressing AstA-R2 in the AstA-AstA-R2 pair seldom expressed TFs (Fig. S11H-I), implying that AstA neuropeptidergic signaling is predominantly regulated by transcriptional control mediated by AstA-R1. However, cells expressing AstA-R2 appear to be less influenced by TF-associated networks. Interestingly, AstA-AstA-R1 pairs are well-documented for their role in modulating developmental maturation (Deveci et al. 2019; Pan and O’Connor 2019), whereas AstA-AstA-R2 signaling is crucial for sleep-deprivation-induced energy wasting in the gut or host survival during intestinal infections (Donlea et al. 2018; Li et al. 2023). Thus, NP-NPR signaling can employ differential transcriptional control to modulate behavior and physiology by distinguishing NPRs in various tissues with distinct associations with TF networks. The AstC-AstC-R pairs display a more nuanced regulation by different NP-NPR pair combinations, revealing that only head tissue cells expressing AstC-R2 are regulated by TF networks, whereas those in the body are not (Fig. S11J-M).

In contrast to the head-biased transcriptional control observed in the AstC-AstC-R2 pair among cells expressing AstC-R2, the sNPF-sNPF-R pair exhibited an opposing pattern of transcriptional control, indicating that cells expressing sNPF-R in body tissue are more susceptible to transcriptional regulation than those in the head (Fig. S11N-O). The Capa-CapaR pair also demonstrated body-biased transcriptional control, similar to the sNPF-sNPF-R pair (Fig. S11P-Q). The NPF-NPFR, FMRFa-FMRFaR, Acp26aAa-SPR, Akh-AkhR, Burs-rk, CCAP-CCAP-R, CNMa-CNMaR, and ETH-ETHR pairs showed typical NPR-biased transcriptional control in both head and body tissues (Fig. S11R-U and Fig. S12A-L). In instances where we detected an absence of TF expression associated with cells expressing NPs or NPRs, we opted for a comprehensive analysis without distinguishing between head and body regions (as illustrated in Fig. S12A-H and Fig. S12K-L).

The neuropeptide CCHamide (CCHa) was initially identified through computational analysis (Nagata 2021). The fruit fly genome encodes two CCHa variants, CCHa1 and CCHa2, which are synthesized by both endocrine cells in the midgut and neurons within the brain. CCHa1, when expressed in the gut, inhibits the transition from sleep to arousal through the modulation of brain dopaminergic neurons and also influences the circadian rhythm regulatory network within the brain (Kuwano et al. 2023; Titos et al. 2023). Conversely, CCHa2 functions as an appetite-stimulating brain-gut peptide, regulating growth by modulating insulin-like peptides in the brain of the fruit fly (Ren et al. 2015; Sano et al. 2015). Our comprehensive analysis suggests that the transcriptional regulation of the CCHa1-CCHa1-R axis is stringent, being mediated by cells expressing CCHa1-R in both head and body tissues. In contrast, for the CCHa2-CCHa2-R axis, it is the cells expressing CCHa2, rather than those expressing CCHa2-R, that are subject to transcriptional control (Fig. S12I-L). Consequently, CCHa1 and CCHa2 employ distinct transcriptional control mechanisms to modulate separate behavioral and physiological responses.

Neuropeptides Diuretic Hormone 44 (Dh44) and Diuretic Hormone 31 (Dh31), which are homologous to mammalian corticotropin-releasing factor (CRF) and calcitonin gene-related peptide (CGRP), respectively, play a pivotal role in the regulation of fluid secretion in *Drosophila*. These peptides, along with their receptors, are implicated in a variety of physiological processes, including the modulation of fluid balance, the sleep-wake cycle, internal nutrient sensing, and CO2-responsive mechanisms (Johnson et al. 2005; Dus et al. 2015; Yang et al. 2018; Lee et al. 2023; Lyu et al. 2023; D.-H. Kim et al. 2024). Notably, the transcriptional regulation of cells expressing receptors for these diuretic hormones is not stringently controlled by transcription factors (Fig. S12Q-T). Diuretic hormones are crucial for maintaining internal homeostasis and regulating innate behaviors, which are vital for the survival of animals. It is intriguing to explore the evolutionary rationale behind the transcriptional control of peptide-expressing cells that do not target receptor-expressing cells. The distinction between innate and adaptive behaviors may provide insights into this divergent system.

While the majority of NP-NPR pairs exhibit either neuropeptide-biased or neuropeptide receptor-biased transcriptional control, the Dsk-CCKLR-17D3 and Gbp5-Lgr1 pairs display a balanced transcriptional control between neuropeptide- and receptor-expressing cells (Fig. S12U-X). It is compelling to investigate the reasons and mechanisms behind these two NP-NPR pairs’ maintenance of unbiased transcriptional control within neuropeptidergic signaling pathways.

Tachykinin (Tk) peptides interact with two distinct receptors, TkR99D and TkR86C, with the latter also being a binding site for neuropeptide natalisin. It has been elucidated that certain modulatory role of Tk within the brain appear to be mediated through TkR99D, while others are through TkR86C. Consequently, Tk signaling exhibits a nuanced and multifaceted pattern of action, which partially intersects with natalisin signaling (Lee et al. 2021; Nässel and Zandawala 2022; Kashio et al. 2023; Wohl et al. 2023). Our comprehensive analysis revealed that cells expressing Tk are subject to more robust TF-associated network regulation compared to cells expressing either TkR99D or TkR86C (Fig. S13A-D). However, when contrasted with natalisin, cells expressing TkR86C exhibit a more stringent transcriptional control in the brain region rather than the body region (Fig. S13E-F). This finding indicates that Tk-expressing neurons are highly regulated by transcriptional mechanisms in the head, whereas neurons expressing natalisin in the head are not. Although natalisin was identified in *Drosophila* and is known to play a role in sexual activity and fecundity, its additional functions remain largely unexplored (Jiang et al. 2013). The variation in the number of TFs associated with Tk, TkR86C, and natalisin suggests that their regulatory mechanisms may vary depending on the context.

Hugin (Hug) neuropeptide engages with two receptors, PK2-R1 and PK2-R2, exerting regulatory effects on developmental timing, systemic growth, and feeding behaviors, with a particular emphasis on hunger-driven actions (Bader et al. 2007; Martelli et al. 2017; Lin et al. 2019; Mizuno et al. 2021; Ohhara et al. 2024). A mutant strain deficient in both PK2-R1 and PK2-R2 exhibited phenotypes similar to the *Hugin* mutant, and developmental aberrations were rescued by the expression of either PK2-R1 or PK2-R2 in PTTH neurons. These findings suggest that Hug-expressing neurons modulate developmental timing and body size through the actions of both PK2-R1- and PK2-R2-expressing neurons (Ohhara et al. 2024). Our detailed analysis reveals a distinctive pattern where the Hugin-PK2-R1 and Hugin-PK2-R2 pairs exhibit Hugin-biased transcriptional control in the head, contrasting with PK2-R1/PK2-R2-biased transcriptional control in the body (Fig. S13G-J). This pattern may indicate that Hugin orchestrates behavior and physiology through both receptors in a head-to-body directional manner. Myosuppressin (Ms) interacts with MsR1 and MsR2 receptors, with MsR1 being implicated in crop enlargement (Hadjieconomou et al. 2020). Intriguingly, our transcription factor network analysis indicates that Ms-MsR1 pairs are predominantly regulated by the transcriptional control of MsR1-expressing cells, whereas MsR2 lacks transcriptional control in head tissue relative to Ms (Fig. S13K-N). Our analysis suggests that MsR1-mediated behavioral modulation is more widespread than that mediated by MsR2.

The *Drosophila melanogaster* species possesses seven insulin-like peptides (ilps), with ilp7 being associated with the receptor Lgr4 and ilp8 with Lgr3 (Garelli et al. 2015; Gontijo and Garelli 2018; Semaniuk et al. 2021; Yeom et al. 2021; Imambocus et al. 2022; Miao et al. 2024). Our analysis has revealed a fascinating dichotomy: the ilp7-Lgr4 pair exhibits a transcriptional control bias towards the neuropeptide receptor (NPR-biased), while the ilp8-Lgr3 pair demonstrates a bias towards the neuropeptide itself (NP-biased) (Fig. S13O-R). This finding aligns with the established importance of Lgr4 in glial cells, influencing both developmental processes and adult behaviors (Imambocus et al. 2022; Miao et al. 2024), and the role of tumor-derived ilp8 in inducing anorexia through the Lgr3 receptor in the brain (Yeom et al. 2021). Our results suggest that these specific insulin-signaling pathways have evolved to employ transcriptional control mechanisms that are either NP-biased or NPR-biased, tailored to their unique physiological roles.

Proctolin (Proc), the inaugural insect neuropeptide to be characterized at the molecular level (Brown 1975; Brown and Starratt 1975; Starratt and Brown 1975; Marder et al. 1986; Orchard et al. 1989; Egerod et al. 2003; Johnson et al. 2003), exerts a direct influence on the body-wall muscles, prompting prolonged, sustained contractions and potentiating contractions evoked by nerve stimulation. Additionally, Proc is known to impact muscle fibers in a cell-selective fashion (Ormerod et al. 2016) through its receptor, ProcR (Johnson et al. 2003). Our intriguing findings reveal that the transcriptional regulation of the Proc-ProcR pair is Proc-biased in the head but shifts to a ProcR-biased pattern in the body (Fig. S13A-B), thereby corroborating the role of Proc in modulating body wall muscle contractions via ProcR (Ormerod et al. 2016). The Nplp1-Gyc76C pair is implicated in the modulation of the innate immune IMD pathway in response to salt stress, particularly within immune and stress-sensing epithelial tissues such as Malpighian tubules and the midgut (Overend et al. 2012). While Nplp1 is known to act through Gyc76C in body tissues, our analysis demonstrates that cells expressing Nplp1 are subject to more robust regulation by the transcription factor-associated network (Fig. S14C-D).

In addition to the aforementioned pairs, other NP-NPR pairs, including Trissin-TrissinR, Pdf-Pdfr, SIFa-SIFaR, Crz-CrzR, Lk-Lkr, and Ptth-Tor, consistently demonstrated that cells expressing NPRs in both the head and body regions are subject to robust transcriptional regulation (Fig. S14E-P). This observation implies that across various NP-NPR pairs, despite the utilization of distinct transcriptional control mechanisms, the cells expressing NPRs are predominantly under stringent transcriptional governance in the majority of cases.

### Empirical validation of transcriptional modulation underpinning NPR-biased regulation of context-dependent behaviors

To empirically validate our analytical findings and the hypothesis that NPR-biased transcriptional mechanisms are a pervasive means of modulating context-dependent behavioral and physiological responses within neuropeptidergic signaling pathways, we undertook quantitative real-time polymerase chain reaction (qRT-PCR) assays. We selected the SIFa-SIFaR interaction for our analysis, given SIFa’s capacity to integrate inputs from a spectrum of neuropeptides and its role in coordinating signaling through a select group of expansive interneurons within neuropeptidergic networks (Martelli et al. 2017; Dreyer et al. 2019; Nässel and Zandawala 2022; W.J. Kim et al. 2024b; T. Zhang et al. 2024b). Additionally, we examined the sNPF-sNPF-R interaction, as sNPF, in conjunction with Tk, exemplifies circuit-specific executive signaling mediated by an array of small-field interneurons (Nässel et al. 2008; Nässel and Zandawala 2022). The Crz-CrzR interaction was included due to its role in hormone-mediated orchestration of signaling, with peptide hormones originating in the brain and corpora cardiaca being released into the circulation to convey state-dependent signals to brain circuits and chemosensory cells in *Drosophila*, thereby influencing sensory processing, behavioral outcomes, and physiological metabolism (Kim et al. 2004; Tayler et al. 2012; Zandawala et al. 2021; Nässel and Zandawala 2022). Lastly, we focused on the AstA-AstA-R1/R2 interaction, as AstA, produced by large neurons that connect several brain regions, exhibits context-specific modulation, albeit with a more limited and focused arborization compared to orchestrating neurons, and is capable of influencing various state-dependent responses such as satiety, hunger, circadian rhythms, and sleep homeostasis across distributed circuits. AstA was selected in part due to its ability to interact with two distinct receptors, highlighting the diverse functions it can mediate (Hergarden et al. 2012; Hentze et al. 2015; Chen et al. 2016; Donlea et al. 2018; Deveci et al. 2019; Wegener and Chen 2022; Li et al. 2023).

For our study, we selected two pivotal environmental contexts—temperature and sexual experiences—that are known to elicit substantial behavioral and physiological shifts in adult *Drosophila*. These contexts serve as critical factors influencing the transcriptional regulation of neuropeptide signaling, thereby modulating complex behaviors in response to environmental cues. Temperature fluctuations are a significant environmental stressor that can profoundly impact gene expression patterns in *Drosophila*, leading to alterations in the transcriptional landscape. Understanding these changes is crucial for deciphering how flies adapt and survive in varying thermal conditions, as it directly influences their physiological responses, metabolic pathways, and behavioral patterns, ultimately determining their fitness and survival in diverse ecological niches (Vazquez et al. 1993; Glaser and Stanewsky 2005; Gibert et al. 2007; MacMillan et al. 2016; Fast and Rosenkranz 2018; Roessingh et al. 2019).

Intriguingly, the mRNA levels of the NPs SIFa, sNPF, and Crz remained unaltered when the environmental temperature was shifted from 22 °C to 29 °C (Fig. 7A-C). In contrast, the mRNA levels of their corresponding NPRs exhibited a significant upregulation following a one-day temperature shift (Fig. 7D-F). This observation implies that the transcriptional regulation of SIFa, sNPF, and Crz in response to temperature changes is predominantly mediated through the modulation of NPR expression levels. Similarly, the mRNA level of AstA was not affected by the temperature shift (Fig. 7G), yet the mRNA level of AstA-R1 was markedly decreased under the same conditions (Fig. 7H), suggesting that the AstA-AstA-R1 interaction plays a role in modulating context-dependent physiological responses by adjusting the transcriptional regulation of NPRs. Notably, the mRNA level of AstA-R2 remained unchanged with temperature variations (Fig. 7I). These findings align with our TF-network analysis, which revealed that AstA-R1, but not AstA-R2, is subject to transcriptional control by TFs (Fig. S11F-I).

**Fig. 7:**
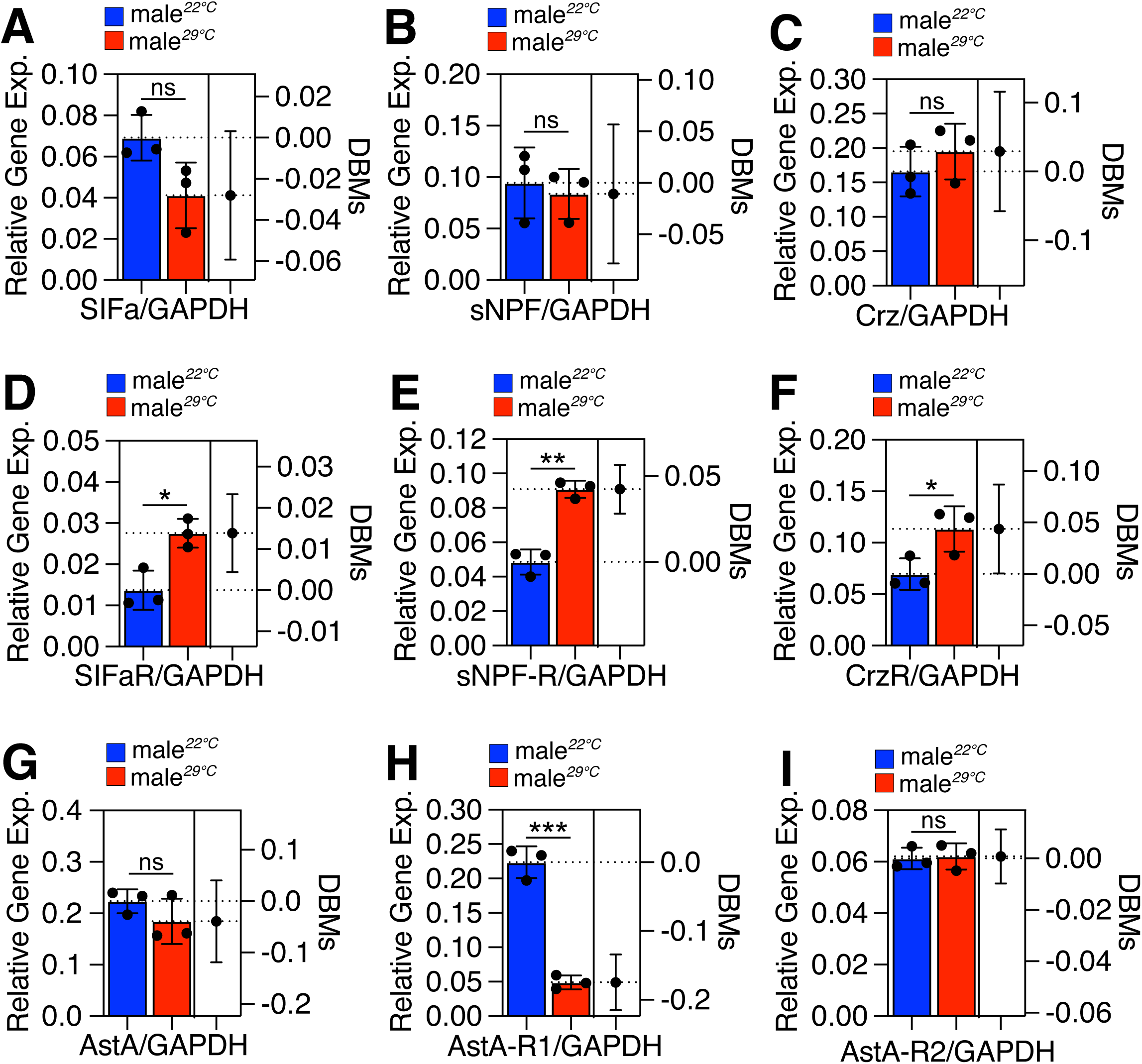
Relative NP and NPR gene expression of Canton-S male flies. **A-I**. Relative expression level of **A**: SIFa, **B**: sNPF, **C**: Crz, **D**: SIFaR, **E**: SIFaR, **F**: CrzR, **G**: AstA, **H**: AstA-R1, **I**: AstA-R2 between Canton-S male flies rearing in 22 and 29 (repeated for 3 times). For detailed methods, see **MATERIALS AND METHODS** section.

Sexual experiences in *Drosophila* are accompanied by significant alterations in the transcriptional landscape, which are essential for the adaptation, survival, and reproductive success of the flies, as these changes underpin the complex behavioral and physiological responses that are crucial for mating and the propagation of their species. Previous studies have established that mating induces comprehensive changes in gene expression profiles in both males and females (McGraw et al. 2004; Mack et al. 2006; Krupp et al. 2008; Ellis and Carney 2010; Gioti et al. 2012; Zhou et al. 2014). In our analysis, the mRNA levels of SIFa and sNPF remained unchanged following sexual experience, whereas the mRNA level of Crz was significantly downregulated (Fig. S15C). Notably, the mRNA levels of SIFaR and CrzR were also significantly reduced by sexual experience, whereas the mRNA level of sNPF-R remained unaffected (Fig. S15D-F). These findings indicate that in the SIFa-SIFaR system, only the NPR levels are transcriptionally modulated by sexual experience, whereas in the Crz-CrzR system, both the NP and NPR levels are transcriptionally regulated by sexual experience. Given the Crz system’s critical role in sexually dimorphic behaviors, these results suggest that both Crz and CrzR may be under transcriptional control to optimize behavioral regulation (Zhao et al. 2010; Tayler et al. 2012; Crickmore and Vosshall 2013; Zer-Krispil et al. 2018; Shao et al. 2019; Thornquist et al. 2020; Oyeyinka et al. 2022). Interestingly, neither the mRNA levels of sNPF nor sNPF-R were altered after sexual experience. However, our previous research identified a critical role for sNPF in modulating male sexual experience-dependent reduced mating behavior (Nässel et al. 2008; Carlsson et al. 2010; W.J. Kim et al. 2016; X. Zhang et al. 2024). This suggests that sNPF signaling may regulate behavior and physiology through alternative modulation mechanisms. Considering that both sNPF and sNPF-R are expressed in multiple neurons and are involved in local transmission, the sNPF system may interact more intimately with the neuromodulatory transmitters system following sexual experience (Nässel et al. 2008; Carlsson et al. 2010).

## DISCUSSION

In the present investigation, we have explored the mechanisms underlying the modulation of context-dependent behaviors by NPs through the transcriptional regulation of their corresponding NPRs. Our approach encompassed a multifaceted analysis, including comparative genomics, assessment of transcription factor networks, and empirical validation, which collectively unveiled a complex regulatory framework. Within this framework, the expression of NPRs emerges as a tightly regulated component, especially in the context of environmental stimuli and physiological changes (Figures 1-7). Our study proposes a hypothesis that redirects attention from NPs to the transcriptional regulation of NPRs, highlighting the significance of NPR gene transcription in modulating context-dependent neuropeptidergic effects on physiological and behavioral outcomes (Figures 1-4). This regulatory mechanism permits a range of responses, from nuanced to pronounced, to a constant NP concentration, thus facilitating organismal adaptation to fluctuating environmental and internal conditions (Figures 5-7). The results of this study contribute to a more nuanced comprehension of neuropeptidergic signaling and offer evolutionary perspectives on the development of complex behavioral and physiological responses in metazoans (Figures 1-7).

Our comprehensive study provides novel insights into the transcriptional regulation of NPRs and its role in modulating context-dependent behaviors and physiological responses. The findings presented herein expand upon the current understanding of neuropeptidergic signaling by revealing the intricate regulatory mechanisms that underpin the multifunctionality of NPs across various organisms, including humans, mice and fruit fly.

### Transcriptional Regulation of NPRs in Behavioral and Physiological Modulation

Our data indicate a significant disparity in the cis-regulatory landscapes of NP and NPR genes, with NPR genes exhibiting a higher number of enhancers, CTCF-binding sites, and open chromatin regions compared to NP genes (Figures 1-4). This suggests a more complex regulatory architecture for NPR genes, which may be crucial for the fine-tuning of neuropeptidergic signaling in response to environmental and physiological cues. The increased length and complexity of NPR genes, as well as their higher propensity for transcriptional modulation, imply a greater capacity for dynamic gene expression and responsiveness (Figures S1C-D). This is further supported by our findings that NPR genes are regulated by more TFs than NP genes, indicating a more intricate regulatory network for NPRs (Figures 6A-I).

### Conservation of Cis-Regulatory Landscapes Across Species

The conservation of the cis-regulatory landscape between human and murine NPR genes underscores the importance of these regulatory elements in the transcriptional control of NP and NPR genes (Figures 1G-L). This conservation may reflect the fundamental role of neuropeptidergic signaling in maintaining homeostasis and adapting to environmental changes across metazoans. The similar quantity of Gene Ontology (GO) annotations for murine NP and NPR genes suggests that both may be involved in a comparable level of functional complexity, despite their distinct biological roles.

### Transcriptional Regulation in Aging and Early Adult Development

Our study also reveals the significance of TF-NPR networks in the aging process of *Drosophila melanogaster*, with NPR expression increasing with age and a higher number of TFs regulating NPRs than NPs (Figures 6A-C). This suggests that NPR genes may be more influenced by key TFs with strong regulatory effects, while NP gene regulation is more dispersed. Furthermore, our analysis of early adult development in *Drosophila* indicates that NP-NPR signaling is highly regulated by TF networks, particularly in a manner biased towards NPR-TF interactions, during the critical period of sexual maturation (Figures S10A-H).

### Differential Transcriptional Regulation of NP-NPR Pairs

Our findings on the differential transcriptional regulation of various NP-NPR pairs provide a nuanced view of how distinct neuropeptidergic signaling pathways may have evolved to employ transcriptional control mechanisms that are either NP-biased or NPR-biased, tailored to their unique physiological roles (Figures S11-S14). This differential regulation may allow for the fine-tuning of complex behaviors and physiological responses in a context-dependent manner.

### Empirical Validation of Transcriptional Modulation

The empirical validation of our analytical findings through qRT-PCR assays further substantiates the role of transcriptional modulation in mediating context-dependent behavioral and physiological responses within neuropeptidergic signaling pathways. The observed changes in mRNA levels of NPRs, but not NPs, in response to temperature shifts and sexual experiences, highlight the importance of transcriptional regulation of NPRs in modulating neuropeptidergic signaling under different environmental and physiological contexts (Figures 7, S15).

### Implications for Understanding Neural Circuit Function and Plasticity

The transcriptional regulation of NPRs, as revealed in this study, has broad implications for our understanding of neural circuit function and plasticity. The ability to modulate NPR expression in response to environmental cues and physiological states allows for the dynamic adjustment of neuropeptidergic signaling, which may underlie the adaptability and complexity of behaviors and physiological processes. This regulatory strategy may be particularly important for survival and reproduction in a changing environment.

In conclusion, our study provides a comprehensive perspective on the transcriptional regulation of NPRs and its role in modulating context-dependent behaviors and physiological responses. The findings not only advance our understanding of neuropeptidergic signaling but also offer insights into the evolution of complex behavioral and physiological responses across metazoans. Future research should focus on elucidating the specific TFs and signaling pathways that regulate NPR expression in different tissues and under varying conditions, as well as exploring the implications of these findings in the context of human health and disease.

## MATERIAL AND METHODS

### Data Collection and Identification

This study utilized publicly available databases and resources to systematically collect data on transcription factor binding sites (TFBS), enhancers, and biological information related to NPs and NPRs in *D. melanogaster*, *M. muscl*e, and *H. Sapiens*.

CTCF-binding site (CTCFBS) annotation data for the human genome were retrieved from the CTCFBSDB 2.0 database (Ziebarth et al. 2012). Based on these annotations, CTCFBSs associated with NP genes and NPR genes were identified and extracted. The number of CTCFBSs for NP and NPR genes was then compared to evaluate their relative abundance.

The human neuropeptide group was obtained from the HGNC Gene Group database (Seal et al. 2022) (https://www.genenames.org/data/genegroup/#!/group/1902). Gene-level details were further supplemented using the Alliance of Genome Resources (Bult and Sternberg 2023) (https://www.alliancegenome.org/gene/). Molecular interaction data, encompassing protein-protein and protein-DNA interactions, were gathered from BioGRID (Oughtred et al. 2021), the IMEx consortium (Porras et al. 2022), WormBase (Sternberg et al. 2024), and FlyBase (Öztürk-Çolak, Marygold, Antonazzo, Attrill, Goutte-Gattat, Jenkins, Matthews, Millburn, Santos, Tabone, Consortium, et al. 2024). These datasets were accessed via the Alliance of Genome Resources, utilizing tab-delimited downloads to enable sorting and filtering based on study-specific criteria.

Mouse tissue expression profiles were obtained from the Gene eXpression Database (GXD) (Baldarelli et al. 2020) provided by The Jackson Laboratory (http://www.informatics.jax.org/expression.shtml).

Data on cis-regulatory modules (CRMs) and other transcriptional regulatory elements were collected from the REDfly database (Kazemian and Halfon 2018; Rivera et al. 2019; Keränen et al. 2022), which is curated information on *Drosophila* regulatory elements.

Single-nucleus transcriptomic data for adult *D. melanogaster* were sourced from the Fly Cell Atlas (FCA) (Li et al. 2022b) This dataset provided crucial insights into tissue-specific and aging-dependent transcriptional landscapes of NPs and NPRs. Processed datasets, crowd-annotation platforms (SCope), and online tools (ASAP) were used to explore tissue-specific expression profiles.

Aging-related transcriptional regulatory patterns were analyzed using the Aging Fly Cell Atlas (AFCA) dataset (Lu et al. 2023b), which offers cellular-resolution insights into the evolution of transcription factor networks regulating NP and NPR genes during aging. These datasets collectively facilitated the investigation of age-dependent molecular mechanisms in *D. melanogaster*.

All datasets were carefully curated, filtered, and validated to ensure consistency and relevance to the study’s objectives.

### Construction of TF-NP/NPRs Regulatory Network

The transcription factor (TF)-NP and NPR regulatory networks were constructed using data from the Fly Cell Atlas (https://flycellatlas.org) (Li et al. 2022b). A total of 17 loom files containing tissue-specific single-nucleus RNA sequencing (snRNA-seq) data were downloaded and processed using the Python package scanpy (Wolf et al. 2018b). The database included information on tissues, genes, cells, gene expression levels, and motif expression levels across cells. Key statistics from the curated data includes Tissues: 17; Genes: 16,373; Cells: 5,078,027; Transcription Factors (TFs): 565; Gene expression records: approximately 450 million. The resulting MySQL database, comprising around 21 GB of data, enables querying of gene expression patterns across the entire *Drosophila melanogaster* dataset, beyond tissue-specific constraints, using SQL commands.

Missing genetic information was supplemented using RNA-Seq RPKM values from the gene_rpkm_matrix_fb_2021_06.tsv file downloaded from FlyBase (https://flybase.org/). These data, originating from FlyAtlas2 (Öztürk-Çolak, Marygold, Antonazzo, Attrill, Goutte-Gattat, Jenkins, Matthews, Millburn, Santos, Tabone, Consortium, et al. 2024), include RNA-Seq and miRNA-Seq measurements across a wide range of tissues, offering valuable insights into gene expression.

Gene lengths were obtained from FlyBase by appending gene symbols to the FlyBase API URL: http://flybase.org/api/sequence/id/+gene_symbol. HTTP requests to this endpoint provided detailed gene information, including lengths.

To construct the TF regulatory network, 10 NP and 13 NPR were selected. In the Fly Cell Atlas, TF-related information is organized using the AUCell algorithm. The relationship between these genes and TFs was analyzed using the following steps: Cells where both NP and NPR are co-expressed were excluded to maximize the distinction between their regulatory patterns. Cells where TFs actively regulate transcription were identified. Information about the genes regulated by TFs within specific tissues was extracted. Gene networks were connected based on cells expressing NPs and the TFs regulating transcription in those cells. Similarly, gene networks were constructed for cells expressing NPRs and their associated TFs. The resulting network graphs allowed for clear visualization of the regulatory differences between TFs involved in NP and NPR expression. The networks were implemented using Python, utilizing the *matplotlib* (Hunter 2007) and *networkx* (github.com/networkx/networkx) libraries for visualization and analysis.

### TF Network Aging Analysis

Single-cell RNA sequencing (scRNA-seq) data from the *Drosophila melanogaster* were obtained from the Aging Fly Cell Atlas website (https://hongjielilab.org/afca/). The data can be easily visualized through a shiny app provided by Hongjie Li et al (Lu et al. 2023a). (https://hongjielilab.shinyapps.io/AFCA/). UMIs data were generated from the h5ad file, comprising a total of 566,254 cells. The Seurat (v4.3.0) package (Hao et al. 2021) was utilized for data analysis. TF regulons and the importance of the pairs at each time point were predicted using SCENIC (Aibar et al. 2017b).

### Code Availability

Code for data cleaning and analysis is provided as part of the replication package. It is available at https://github.com/sulitech/Drosophila- for review. It will be uploaded to the [JOURNAL REPOSITORY] once the paper has been conditionally accepted.

### Other Resources

Anatomical illustrations for internal and sensory organs of *Drosophila* were sourced from (Bilder et al. 2021) and (Lorimer et al. 2014), respectively.

### Fly stock and husbandry

*Drosophila melanogaster* Canton-S line was raised on cornmeal-yeast medium at similar densities to yield adults with similar body sizes. Flies were kept in 12 h light: 12 h dark cycles (LD) at 25 (ZT 0 is the beginning of the light phase, ZT12 beginning of the dark phase). DF (here after) genetic modified fly line in this study, which harbors a deletion of a specific genomic region that includes the sex peptide receptor (SPR) (Parks et al. 2004; Yapici et al. 2008). Previous studies have demonstrated that virgin females of this line exhibit increased receptivity to males (Yapici et al. 2008). For naïve CS males, 40 males from the same strain were placed into a vial with food for 5 days, and moved to 29u before RNA extraction assay. For experienced males, 40 males from the same strain were placed into a vial with food for 6 days and twice as number DF virgin females were introduced into vials for last 1 day before assay as introduced in previous study (Sun et al. 2024).

### RNA extraction and cDNA synthesis

RNA was extracted from 50 preparations of 6-day-old Canton-S males using the RNA isolation kit (Vazyme), following the manufacturer’s protocol. For each condition, flies were reared in 22 and 29 respectively for one day, and virgin female flies in doubled number were introduced to vials for one day to give male flies sexual experience. And first-strand cDNA was synthesized from 1μg of RNA template with random primers using SPARK script - RT plus kit (SparkJade).

### Quantitative RT-PCR

The expression levels of SIFa, sNPF, Crz, SIFaR, sNPF-R, CrzR, AstA, AstA-R1, AstA-R2 in Canton-S male flies in different conditions were analyzed by quantitative real-time RT-PCR with SYBR Green qPCR MasterMix kit (Selleckchem). The primers of RT-PCR are SIFa, F: 5’- ACTGCAAGATGGCTCTTCG -3’; R: 5’-CGGCATTTCCACATTCAGTC -3’; sNPF, F: 5’- GTGTTCCTCAGTTCGAGGCAA-3’; R: 5’-AGTTCAAAAGCGAGTTGTACCA-3’; Crz, F: 5’-GAAACTGTGTCCCCGGTTCG-3’; R: 5’-AGAGTTGCTCAGTCTGGGATG-3’; SIFaR, F: 5’-GGGCACGGTTCTCACTACG-3’; R: 5’-CGATAGGTTCAGCAAGTTGGAA-3’; sNPF-R, F: 5’-AACTGGTTGTGAATGATCCCG-3’; R: 5’-CCAACTGGAGCCTAACGTCG-3’; CrzR, F: 5’-AATCCGGACAAAAGGCTGGG -3’; R: 5’-AGGTGGAAGGCACCGTAGAT-3’; AstA, F: 5’-TCCCTTCACGCCCACCTCCT-3’; R: 5’-TACCGCTCCACCCGCTTGTC-3’; AstA-R1, F: 5’-CGCAGAGTCACGAAAGGG-3’; R: 5’-CGCCACCACATGGGATAT-3’; AstA-R2, F:5’-CCAACCTGATGATTGTCAATCTGGC-3’; R: 5’- GGTAATGTTCTCCGTCCTCATCATC-3’.qPCR reactions were performed in triplicate, and the specificity of each reaction was evaluated by dissociation curve analysis. Each experiment was replicated three times. PCR results were recorded as threshold cycle numbers (Ct). The fold change in the target gene expression, normalized to the expression of internal control gene (GAPDH) and relative to the expression at time point 0, was calculated using the 2 −ΔΔCT method as previously described (Livak and Schmittgen 2001). The results are presented as the mean ± SD of three independent experiments.

### Statistical Tests

When dataset passed test for normal distribution (Kolmogorov-Smirnov tests, p > 0.05), we used two-sided Student’s t tests. The mean ± standard error (s.e.m) (*\*\*\*\* = p < 0.0001, *** = p < 0.001, ** = p < 0.01, * = p < 0.05*). All analysis was done in GraphPad (Prism). Individual tests and significance are detailed in figure or figure legends.

Besides traditional *t*-test for statistical analysis, we added estimation statistics for all analysis in two group comparing graphs. In short, ‘estimation statistics’ is a simple framework that—while avoiding the pitfalls of significance testing—uses familiar statistical concepts: means, mean differences, and error bars. More importantly, it focuses on the effect size of one’s experiment/intervention, as opposed to significance testing (Claridge-Chang and Assam 2016). In comparison to typical NHST plots, estimation graphics have the following five significant advantages such as (1) avoid false dichotomy, (2) display all observed values (3) visualize estimate precision (4) show mean difference distribution. And most importantly (5) by focusing attention on an effect size, the difference diagram encourages quantitative reasoning about the system under study (Ho et al. 2019). Thus, we conducted a reanalysis of all our two group data sets using both standard *t* tests and estimate statistics. In 2019, the Society for Neuroscience journal eNeuro instituted a policy recommending the use of estimation graphics as the preferred method for data presentation (Bernard 2021).

For datasets that did not exhibit normal distribution, as determined by the Kolmogorov-Smirnov test, we employed non-parametric statistical methods to analyze the data. Specifically, we utilized the Mann-Whitney U test to compare the distributions between two independent groups (Whitfield and Siegel 1957). This test is appropriate for ordinal data and does not assume a normal distribution, making it a robust choice for our analysis. The results of these comparisons were visualized using rank plots, which provide a graphical representation of the data distribution and the test statistics. An example of such visualization is presented in Figures 4B-C (Conover and Iman 1981).

## FIGURE LEGEND

**Figure S1. Comparative analysis of cis-regulatory elements and gene characteristics between NP and NPR genes across species.**

**A-B**. Average number of transcription factor binding sites (TFBSs) per gene in *H. sapiens*, comparing NP and NPR genes.

**C-D**. Gene length distributions in *M. musculus* NP and NPR genes.

**E-F**. Mean length of transcription factor binding site hotspot areas (TFBS-HSA) in *D. melanogaster* NP and NPR genes.

**G-H**. Number of cis-regulatory modules (CRMs) identified in *Drosophila* NP (G) and NPR (H) genes, indicating variation in regulatory complexity between the two gene types.

**I-J**. Mean length of Cis-regulatory modules (CRMs) in *D. melanogaster* NP and NPR genes.

**K-L**. Tissue-specific expression patterns of NP and NPR genes in *D. melanogaster*.

**Figure S2. Comparative analysis of gene ontology (GO) annotations and transcription factor binding sites (TFBS) between NP and NPR genes in mouse.**

**A-B**. Number of GO annotations for NP and NPR genes in mouse, both unpaired and paired.

**C-D**. Number of TFBSs identified by GXD tissue expression analysis for NP and NPR genes in mouse, both unpaired and paired.

**Figure S3. Spatial Expression Profiles of NPs and NPRs in across *D. melanogaster* Tissues. A-Q**. Scatter Plot of Total Gene Expression (TPM) and Total Cell Count (TCC) by Tissues in *D. melanogaster*, including Head (a), Body (b), Antenna (c), Body Wall (d), Fat Body (e), Gut (f), Haltere (g), Heart (h), Leg (i), Male Reproductive Glands (j), Malpighian Tubule (k), Oenocyte (l), Ovary (m), Proboscis (n), Testis (o), Trachea (p), Wing (q). Dots representing NP or NPR genes are color-coded by cell count (C. C) threshold. The log scale on the y-axis indicates the transformation applied to the TPM values for visualization purposes.

**Fig. S4. Transcription factors (TFs) that specifically regulate NP and NPR in various tissues.**

**A**: TF-NP and TF-NPR regulatory networks across all tissues (see Fig. 4A).

**B**: Total number of TFs uniquely regulating NP and NPR genes across all tissues.

**C**: Total number of TF motifs uniquely regulating NP and NPR genes across all tissues.

**D**: Total number of TF motifs uniquely regulating NP and NPR genes in specific tissues.

**E**: TF regulatory network for NPs and NPRs in the head region.

**F**: Number of TFs specifically regulating NP and NPR genes in the head.

**G**: TF regulatory network for NPs and NPRs in the body region.

**H**: Number of TFs specifically regulating NP and NPR genes in the body.

**I**: Comparison of TF counts regulating NP and NPR genes across other tissues, including the gut, fat body, body wall, heart, oenocyte, and wing.

**Fig. S5. Tissue-specific TFs that regulate NPs and NPRs**

**A**: Network of TFs that regulate NPs and NPRs in the Fat Body (see Fig. 4A).

**B**: Number of TFs regulating NPs and NPRs in the Fat Body.

**C**: Number of NPs and NPRs regulated by TFs in the Fat Body.

**D**: Network of TFs that regulate NPs and NPRs in the Gut.

**E**: Number of TFs that regulate NPs and NPRs in the Gut. **F**: Number of NPs and NPRs regulated by TFs in the Gut.

**G**: Network of TFs that regulate NPs and NPRs in the Heart.

**H**: Number of TFs that regulate NPs and NPRs in the Heart.

**I:** Number of NPs and NPRs regulated by TFs in the Heart.

**J:** Network of TFs that regulate NPs and NPRs in the Malpighian tubules.

**K**: Number of TFs that regulate NPs and NPRs in the Malpighian tubules.

**L**: Number of NPs and NPRs regulated by TFs Malpighian tubules.

**M**: Network of TFs that regulate NPs and NPRs in the Oenocyte.

**N**: Number of TFs that regulate NPs and NPRs in the Oenocyte.

**O**: Number of NPs and NPRs regulated by TFs in the Oenocyte.

**P**: Network of TFs that regulate NPs and NPRs in the Trachea.

**Q**: Number of TFs that regulate NPs and NPRs in the Trachea.

**R**: Number of NPs and NPRs regulated by TFs in the Trachea.

**Fig. S6. Tissue-specific TFs that regulate NPs and NPRs**

**A**: Network of TFs that regulate NPs and NPRs in the Antenna (see Fig. 4A).

**B**: Number of TFs that regulate NPs and NPRs in the Antenna.

**C**: Number of NPs and NPRs regulated by TFs in the Antenna.

**D**: Network of TFs that regulate NPs and NPRs in the Body wall.

**E**: Number of TFs that regulate NPs and NPRs in the Body wall.

**F**: Number of NPs and NPRs regulated by TFs in the Body wall.

**G**: Network of TFs that regulate NPs and NPRs in the Haltere.

**H**: Number of TFs that regulate NPs and NPRs in the Haltere.

**I**: Number of NPs and NPRs regulated by TFs in the Haltere.

**J**: Network of TFs that regulate NPs and NPRs in the Leg.

**K**: Number of TFs that regulate NPs and NPRs in the Leg.

**L**: Number of NPs and NPRs regulated by TFs Leg.

**M**: Network of TFs that regulate NPs and NPRs in the Proboscis.

**N**: Number of TFs that regulate NPs and NPRs in the Proboscis.

**O**: Number of NPs and NPRs regulated by TFs in the Proboscis.

**P**: Network of TFs that regulate NPs and NPRs in the Wing.

**Q**: Number of TFs that regulate NPs and NPRs in the Wing.

**R**: Number of NPs and NPRs regulated by TFs in the Wing.

**Fig. S7. Tissue-specific TFs that regulate NPs and NPRs**

**A**: Network of TFs that regulate NPs and NPRs in the Male reproductive glands (see Fig. 4A).

**B**: Number of TFs that regulate NPs and NPRs in the Male reproductive glands.

**C**: Number of NPs and NPRs regulated by TFs in the Male reproductive glands.

**D**: Network of TFs that regulate NPs and NPRs in the Ovary.

**E**: Number of TFs that regulate NPs and NPRs in the Ovary.

**F**: Number of NPs and NPRs regulated by TFs in the Ovary.

**G**: Network of TFs that regulate NPs and NPRs in the Testis.

**H**: Number of TFs that regulate NPs and NPRs in the Testis.

**I**: Number of NPs and NPRs regulated by TFs in the Testis.

**Fig. S8. TFs that regulate NPs and NPRs in all tissues of males and females.**

**A**: Network of TFs that regulate NPs and NPRs in all tissues of males (see Fig. 4A).

**B**: Number of TFs that regulate NPs and NPRs in all tissues of males.

**C**: Number of NPs and NPRs regulated by TFs in all tissues of males.

**D**: Network of TFs that regulate NPs and NPRs in all tissues of females.

**E**: Number of TFs that regulate NPs and NPRs in all tissues of females.

**F**: Number of NPs and NPRs regulated by TFs in all tissues of females.

**Fig. S9. TFs that regulate NPs and NPRs in all tissues**

**A**: Network of TFs that regulate **NTs** and **NTRs** in all tissues (see Fig. 4A).

**B**: Number of TFs that regulate **NTs** and **NTRs** in all tissues.

**C**: Number of **NTs** and **NTRs** regulated by TFs in all tissues.

**D**: Network of TFs that regulate **NTs** and **NTRs** in the Head.

**E**: Number of TFs that regulate **NTs** and **NTRs** in the Head.

**F**: Number of **NTs** and **NTRs** regulated by TFs in the Head.

**G**: Network of TFs that regulate **NTs** and **NTRs** in the Body.

**H**: Number of TFs that regulate **NTs** and **NTRs** in the Body.

**I:** Number of **NTs** and **NTRs** regulated by TFs in the Body.

**Fig. S10. TFs that regulate NPs and NPRs in all tissues by age**

**A**: Cell counts of NPs and NPRs expressed by age.

**B**: Network of TFs that regulate NPs and NPRs in all tissues on Day 1 (see Fig. 4A).

**C**: Number of TFs that regulate NPs and NPRs in all tissues on Day 1.

**D**: Number of NPs and NPRs regulated by TFs in all tissues on Day 1.

**E**: Network of TFs that regulate NPs and NPRs in all tissues on Day 3.

**F**: Number of TFs that regulate NPs and NPRs in all tissues on Day 3.

**G**: Number of NPs and NPRs regulated by TFs in all tissues on Day 3.

**H**: Number of TFs remaining after removing those expressed on Day 3 from Day 5, and those expressed on Day 1 from Day 3.

**Figure S11: Comparative Analysis of TF Networks in ilp/InR-Expressing Cells and NP-NPR Pairs Across Head and Body Regions**

**A-E**. Regulatory TF Expression in ilp- and InR-Expressing Cells. (a) Quantification of TF- regulated cell counts in ilp-positive and InR-positive cells across total, head, and body regions. (b and d) Network maps illustrating TF interactions in ilp and InR cellular networks for the head (b) and body (d) region. (c and e) Violin plots showing the distribution of TF expression in ilp-and InR-expressing cells among head (c) and body (e) region.

**F-U**. Differences in regulatory TF expression between head and body across NP-NPR pairs: (f-g) AstA–AstA-R1, (h-i) AstA–AstA-R2, (j-k) AstC–AstC-R1, (l-m) AstC–AstC-R2, (n-o) sNPF– sNPF-R, (p-q) Capa–CapaR, (r-s) NPF–NPFR, and (t-u) FMRFa–FMRFaR.

**Figure S12: Comparative Analysis of TF Networks in ilp/InR-Expressing Cells and NP- NPR Pairs Across Head and Body Regions.**

**A-X**. Differences in regulatory TF expression between head and body across NP–NPR pairs: (a-b) Acp26aAa–SPR, (c-d) Akh–AkhR, (e-f) Burs–rk, (g-h) CCAP–CCAP-R, (i-j) CNMa–CNMaR, (k-l) ETH–ETHR, (m-n) CCHa1–CCHa1-R, (o-p) CCHa2–CCHa2-R, (q-r) Dh31–hec, (s-t) Dh44–Dh44-R1, (u-v) Dsk–CCKLR-17D3, and (w-x) Gbp5–Lgr1.

**Figure S13: Continuation of Figure S12.**

**A-Q**. Differences in regulatory TF expression between head and body across NP–NPR pairs: (a-b) Tk–TkR99D, (c-d) Tk–TkR86C, (e-f) natalisin–TkR86C, (g-h) Hug–PK2-R1, (i-j) Hug–PK2-R2, (k-l) Ms–MsR1, (m-n) Ms–MsR2, (o-p) ilp7–Lgr4, and (q-r) ilp8–Lgr3.

**Figure S14: Continuation of Figure S13.**

**A-P**. Differences in regulatory TF expression between head and body across NP–NPR pairs: (a-b) Proc–ProcR, (c-d) Nplp1–Gyc76C, (e-f) Trissin–TrissinR, (g-h) Pdf–Pdfr, (i-j) SIFa–SIFaR, (k-l) Crz–CrzR, (m-n) Lk–Lkr, and (o-p) Ptth–Tor.

Fig. S15: Relative gene expression of Canton-S male flies.

**A-I**. Relative expression level of **A**: SIFa, **B**: sNPF, **C**: Crz, **D**: SIFaR, **E**: SIFaR, **F**: CrzR, **G**: AstA, **H**: AstA-R1, **I**: AstA-R2 between Canton-S male flies rearing in 29 sexual experience (repeated for 3 times).

## ACKNOWLEDGEMENTS

We express our sincere appreciation to the flySCope platform for providing invaluable resources that facilitated the analysis and validation of our hypothesis. The genetic research conducted within this study was significantly enhanced by the comprehensive resources offered by the FlyBase website. We extend our gratitude to the FlyBase team for their diligent maintenance of this extensive *Drosophila* database (Pierre and McQuilton 2009; Attrill et al. 2016; Gramates et al. 2022; Jenkins et al. 2022; Öztürk-Çolak, Marygold, Antonazzo, Attrill, Goutte-Gattat, Jenkins, Matthews, Millburn, Santos, Tabone, Perrimon, et al. 2024). This research was financially supported by the Startup funds from the HIT Center for Life Science, awarded to WJK. It is important to note that the funding bodies played no role in the study design, data collection and analysis, decision to publish, or the preparation of the manuscript.

## ETHICS APPROVAL AND STATEMENT OF ANIMAL RESEARCH COMPLIANCE

All animal experiments reported in this manuscript were conducted in compliance with the ARRIVE guidelines and adhered to the U.K. Animals (Scientific Procedures) Act, 1986 and associated guidelines, EU Directive 2010/63/EU for animal experiments, or the National Research Council’s Guide for the Care and Use of Laboratory Animals (Animals et al. 2011).

## CONSENT TO PARTICIPATE

The research described in this paper does not involve any human participants. Therefore, no consent to participate was obtained.

## CONSENT FOR PUBLICATION

All authors have given consent for the publication of this work.

## AUTHOR CONTRIBUTIONS

**Conceptualization:** WJK and SHR.

**Data curation:** WJK, SHR, and ZW.

**Formal analysis:** WJK, SHR, ZW, and YW.

**Funding acquisition:** WJK.

**Investigation:** WJK.

**Methodology:** WJK, SHR, and ZW.

**Project administration:** WJK. **Resources:** WJK and SHR.

**Supervision:** WJK and DHL.

**Validation:** WJK, SHR, ZW, and YW.

**Visualization:** WJK, SHR, ZW, and YW.

**Writing – original draft:** WJK, SHR, ZW, and YW.

**Writing – review & editing:** WJK, SHR, ZW, and YW.

## DATA AVAILABILIITY STATEMENT

All data and reagents reported in this paper will be shared by the corresponding author upon request.

## DECLARATION OF GENERATIVE AI AND AI-ASSISTED TECHNOLOGIES IN THE WRITING PROCESS

During the creation of this work, the author(s) utilized ChatGLM (https://chatglm.cn/) and KIMI (https://kimi.moonshot.cn/) to rephrase English sentences, verify English grammar, and detect plagiarism, as none of the authors of this paper are native English speakers. After using this tool/service, the author(s) reviewed and edited the content as needed and take(s) full responsibility for the content of the publication.

## STATEMENT OF ANIMAL RESEARCH COMPLIANCE

All animal experiments reported in this manuscript were conducted in compliance with the ARRIVE guidelines and adhered to the U.K. Animals (Scientific Procedures) Act, 1986 and associated guidelines, EU Directive 2010/63/EU for animal experiments, or the National Research Council’s Guide for the Care and Use of Laboratory Animals.

## CODE AVAILABILITY

All code is available upon request.

## DATA AVAILABILITY STATEMENT

Strains are available upon request. The authors affirm that all data necessary for confirming the conclusions of the article are present within the article, figures, and tables.

